# *In vivo* expression of engineered IsPETase and LCC variants in *Drosophila melanogaster* reveals host tolerance and differential catalytic activity

**DOI:** 10.64898/2026.06.04.729872

**Authors:** Valentina Pirillo, Federica Barca, Daniele Bruno, Sara Caramella, Claudia Fontana, Matteo Battistolli, Elena Catelan-Carphio, Davis Roma, Morena Casartelli, Silvia Caccia, Alessandro Grapputo, Gianluca Tettamanti, Gianluca Molla, Federica Sandrelli

**Author notes:** These authors contributed equally: Valentina Pirillo, Federica Barca. **Co-corresponding authors** Gianluca Molla email address; Federica Sandrelli.

## Abstract

Insects offer promising opportunities for organic waste bioconversion; however, they cannot efficiently degrade synthetic polymers such as polyethylene terephthalate (PET). Here, we generated transgenic *Drosophila melanogaster* lines to express *in vitro*-evolved variants of two PET-degrading enzymes with distinct biochemical properties: an engineered *Ideonella sakaiensis* PETase variant (TS-ΔIsPET) and a leaf-branch compost cutinase variant (TA-ΔLCC). Both enzymes, fused to a *Drosophila* gut-derived secretory signal, were produced and secreted by both *Drosophila* cultured S2R^+^ cells and transgenic larvae. Both enzymes were glycosylated upon secretion, a post-translational modification that did not abolish their catalytic activity. Notably, TA-ΔLCC displayed ∼6-fold higher esterase activity than TS-ΔIsPET in larval extracts and TA-ΔLCC-containing extracts depolymerised PET nanoparticles *in vitro* under enzyme-favourable conditions. Transgenic flies showed normal development, fertility and survival. Morphological and biochemical analysis confirmed that TA-ΔLCC expression did not alter midgut structure and function. Together, these results establish *Drosophila melanogaster* as a model for functional expression and comparative evaluation of engineered PET-degrading enzymes and identify TA-ΔLCC as a promising candidate for exploitation in insect species relevant to plastic contaminated waste bioconversion.

**Highlights:**

- Transgenic *D. melanogaster* enables *in vivo* study of engineered PET enzymes
- Engineered TS-ΔIsPET and TA-ΔLCC are functional in larval extracts
- TA-ΔLCC was selected for PET nanoparticle assays due to higher pNPA activity
- *D. melanogaster* model enables comparative evaluation of PET-degrading enzymes

## 1. Introduction

Plastics have become an integral part of our daily life due to their low cost, versatility, and durability, progressively replacing other materials in many applications [1]. As plastic production continues to rise, multiscale commitments have emerged to limit plastic emissions into the environment. In this context, reducing plastic pollution has been included into the United Nations Sustainable Development Goals [2,3]. Over 70 % of produced plastic ultimately becomes waste, with approximately 47 % consisting of plastic packaging. This includes polyethylene terephthalate (PET), a thermoplastic polymer synthesised from terephthalic acid (TPA) and ethylene glycol (EG). PET is widely used in packaging and textile fibre manufacturing, with an annual global production exceeding 70 million tons [4], representing nearly 16 % of European plastic consumption in the packaging industry and ∼12 % of global plastic solid waste [5–7].

Collection systems and recycling procedures for PET, particularly in the form of bottles, are well-established and efficient. However, other PET fractions, such as those originating from consumer packaging materials, often escape established collection streams. In this context, plastics contaminate the organic fraction of municipal solid waste (OFMSW), that can represent up to 57 % of the total waste generated in low-income countries [8,9].

OFMSW is mainly processed through anaerobic digestion and composting, generating materials used as soil conditioners or fertilisers [10]. However, plastic contaminants in these streams can be redistributed onto crops and agricultural products, affecting plant performance, soil quality, and soil-dwelling organisms [11,12]. Furthermore, these contaminants may infiltrate groundwater and aquatic ecosystems [13]. Therefore, addressing this issue is critical for reducing the environmental impact of post-consumer PET, thereby advancing waste management sustainability [14].

Insect larvae, such as those of the coleopterans *Tenebrio molitor* (mealworm) and *Zophobas morio* (superworm), and the dipteran *Hermetia illucens* (Black Soldier Fly, BSF), have been proposed for the treatment of different organic waste streams [15–18]. Specifically, BSF shows a high metabolic flexibility and bioconversion efficiency, enabling the valorisation of OFMSW into valuable biomolecules, including proteins, lipids and chitin [15,16,19].

Beyond organic waste recycling, various insect species [e.g., the lepidopterans *Galleria mellonella* (greater wax moth) and *Plodia interpunctella* (Indian meal moth), the mealworm *T. molitor*, the superworms, and, to a more limited extent, BSF] have been reported to grow in the presence of different plastics and promote their partial degradation [20–30]. However, the mechanisms underlying this capability are still under investigation. Current evidence suggests that plastic modification results from a combination of insect chewing activity and biodegradation processes, mainly driven by gut microorganisms or, in some cases, by insect-produced enzymes [21–28,30,31]. Nevertheless, degradation efficiency is highly variable and depends on polymer properties and experimental conditions [24,28]. In addition, the fate of plastic-derived residues remains poorly characterised. Thus, while insect-associated plastic degradation represents an intriguing biological phenomenon, significant challenges remain for its translation into practical plastic waste treatment strategies [24,28].

In the last two decades, extensive research has focused on microbial enzymes able to depolymerise PET [32]. Among these, the most studied are the cutinase isolated from a leaf-branch compost (LCC) [33] and the PET hydrolase from the mesophilic bacterium *Ideonella sakaiensis* (IsPETase) [34,35]. Here, we maintain the term IsPETase, although the organism has been recently reclassified as *Piscinibacter sakaiensis* [36]. Since the depolymerisation efficiency and stability of wild-type enzymes are inadequate for industrial-scale applications [37], extensive research has been dedicated to developing engineered forms of both IsPETase and LCC with improved catalytic properties [32,38–40]. Recent advances in genome editing have enabled the generation of genetically modified insects expressing engineered plastic-degrading enzymes of microbial origin in their intestinal tract, potentially improving their capacity to process plastic-contaminated waste.

To explore this approach, the dipteran *Drosophila melanogaster* (the fruit fly) represents a powerful model for studying heterologous enzymes in an insect host. Consistently, the expression of an industrially relevant fungal laccase has been demonstrated in this system [41].

Although *D. melanogaster* is not known to degrade plastic materials, its genetic tools enable controlled *in vivo* studies of plastic-degrading enzymes [42]. In this context, a recent work has demonstrated the functional expression of wild-type PETase in *D. melanogaster* [43], supporting its use as a platform for the evaluation of engineered PET-degrading enzymes. Using this model, we assessed the expression and functional activity of two engineered variants of IsPETase and LCC (TS-ΔIsPET and TA-ΔLCC, respectively), designed to efficiently depolymerise PET at moderate temperatures. Our results show that transgenic *D. melanogaster* allows the functional expression of these enzymes, enabling their comparative evaluation in an insect host. In addition, by extending this approach to an engineered LCC variant not previously explored in this model, we identify TA-ΔLCC as a candidate for further investigation in insect-based strategies addressing PET-contaminated waste.

## 2. Results

### 2.1. *In vitro* evolution of PET-degrading enzymes active at moderate temperatures

The PET hydrolase from *I. sakaiensis* (IsPETase) and the leaf-branch compost cutinase (LCC) are two promising PET hydrolysing enzymes for biotechnological applications. We previously evolved a thermostable variant of IsPETase, TS-ΔIsPET (Fig. 1A), that shows a 1.8-fold higher activity on PET nanoparticles than the wild-type at 30 °C (Fig. 1C) [44]. LCC is a thermostable protein with a melting temperature (T_m_) of 80.8 °C, higher than TS-ΔIsPET (55.3 °C) [45]. LCC exhibits a higher efficiency toward semi-crystalline PET at elevated temperatures, whereas IsPETase preferentially degrades amorphous PET under mesophilic conditions. TS-ΔIsPET produces a mixture of mono(hydroxyethyl)terephthalate (MHET) and TPA from PET depolymerisation, LCC produces almost only TPA [46].

**Fig. 1.**
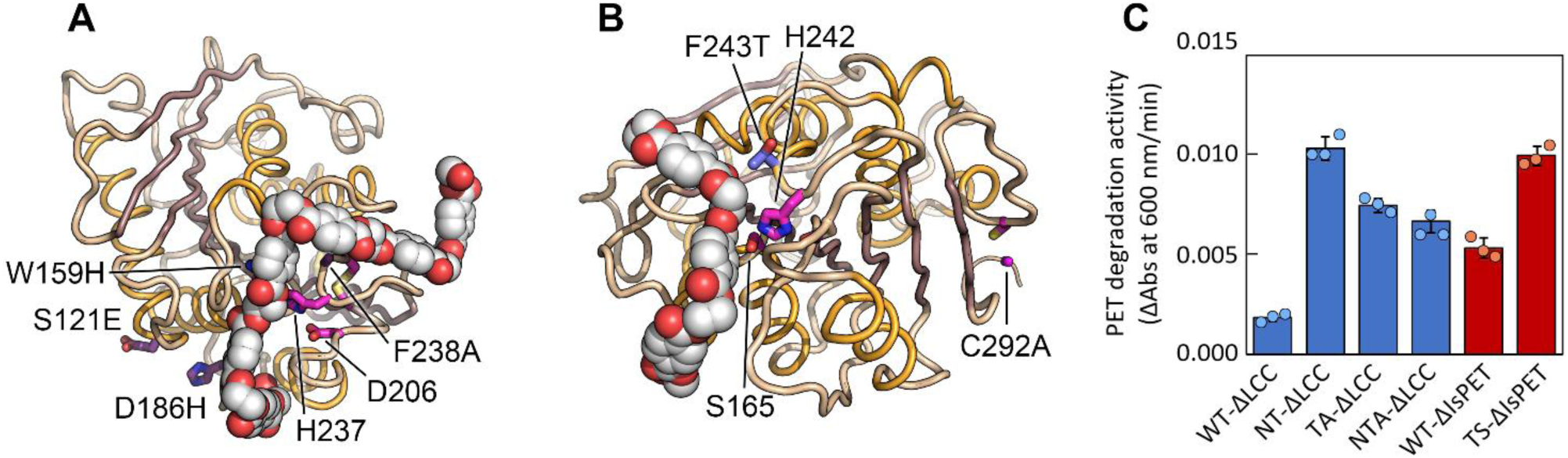
Identification of TS-ΔIsPET and TA-ΔLCC variants for expression in the *Drosophila melanogaster* model. **A, B** 3D structural models of TS-ΔIsPET (**A**) and TA-ΔLCC (**B**) variants. **C** Relative activity (mean ± SEM) of IsPETase and ΔLCC variants on PET nanoparticles (0.1 mg/mL) at 30 °C. Reaction buffer: 25 mM Tris-HCl, 200 mM NaCl, pH 8.0. ΔAbs/min values are shown as absolute values. For each variant, 3 replicates (dots) were analysed.

To increase ΔLCC activity at lower temperatures, we removed the C-terminal disulfide bridge between cysteines C275 and C292 by site-directed mutagenesis. This modification is expected to enhance the structural flexibility of the protein, reducing its thermal stability and increasing its activity at 30 °C [33]. The C292A substitution was introduced into previously improved single F243T-ΔLCC and double S101N/F243T-ΔLCC (NT-ΔLCC) variants developed in our laboratory [44,45], producing the double (F243T/C292A-ΔLCC; TA-ΔLCC) and the triple (S101N/F243T/C292A-ΔLCC; NTA-ΔLCC) variants, respectively. TS-ΔIsPET and TA-ΔLCC were selected for expression in *D. melanogaster* (Fig. 1A, B). Specifically, TA-ΔLCC was chosen despite showing a marginal lower activity on PET nanoparticles at 30 °C compared to the NT-ΔLCC variant, as it lacks disulfide bridges (Fig. 1C), thus increasing the diversity of the selected biocatalysts (TS-ΔIsPET possesses two disulfide bridges). TA-ΔLCC retained a significant activity at moderate temperature (e.g., the activity at 37 °C was still ∼41 % compared to the activity at 55 °C) (Supplementary Fig. 1). The T_m_ of TA-ΔLCC was 69.9 ± 0.1, ∼11 °C lower compared to the wild-type ΔLCC (Supplementary Fig. 2) [45].

### 2.2. Expression of *in vitro-*evolved PET-degrading enzymes in *D. melanogaster* cell lines

To enable active secretion of the *in vitro*-evolved PET-degrading enzymes in *D. melanogaster*, we designed the Dm-TS-ΔIsPET and Dm-TA-ΔLCC chimeric proteins. Their sequences include an N-terminal secretory signal peptide (SP) derived from a *D. melanogaster* chymotrypsin-like protein highly expressed in the midgut, followed by either the TS-ΔIsPET or TA-ΔLCC variants, and a C-terminal His tag (Fig. 2A and Supplementary Fig. 3). The synthesis and secretion of these recombinant PET hydrolytic enzymes were first evaluated *in vitro,* generating stable S2R+ *D. melanogaster* cell lines, carrying either the *Dm-TS-ΔIsPET* or *Dm-TA-ΔLCC* DNA sequences cloned into the pAc5-STABLE2-neo expression vector (Supplementary Fig. 4).

**Fig. 2.**
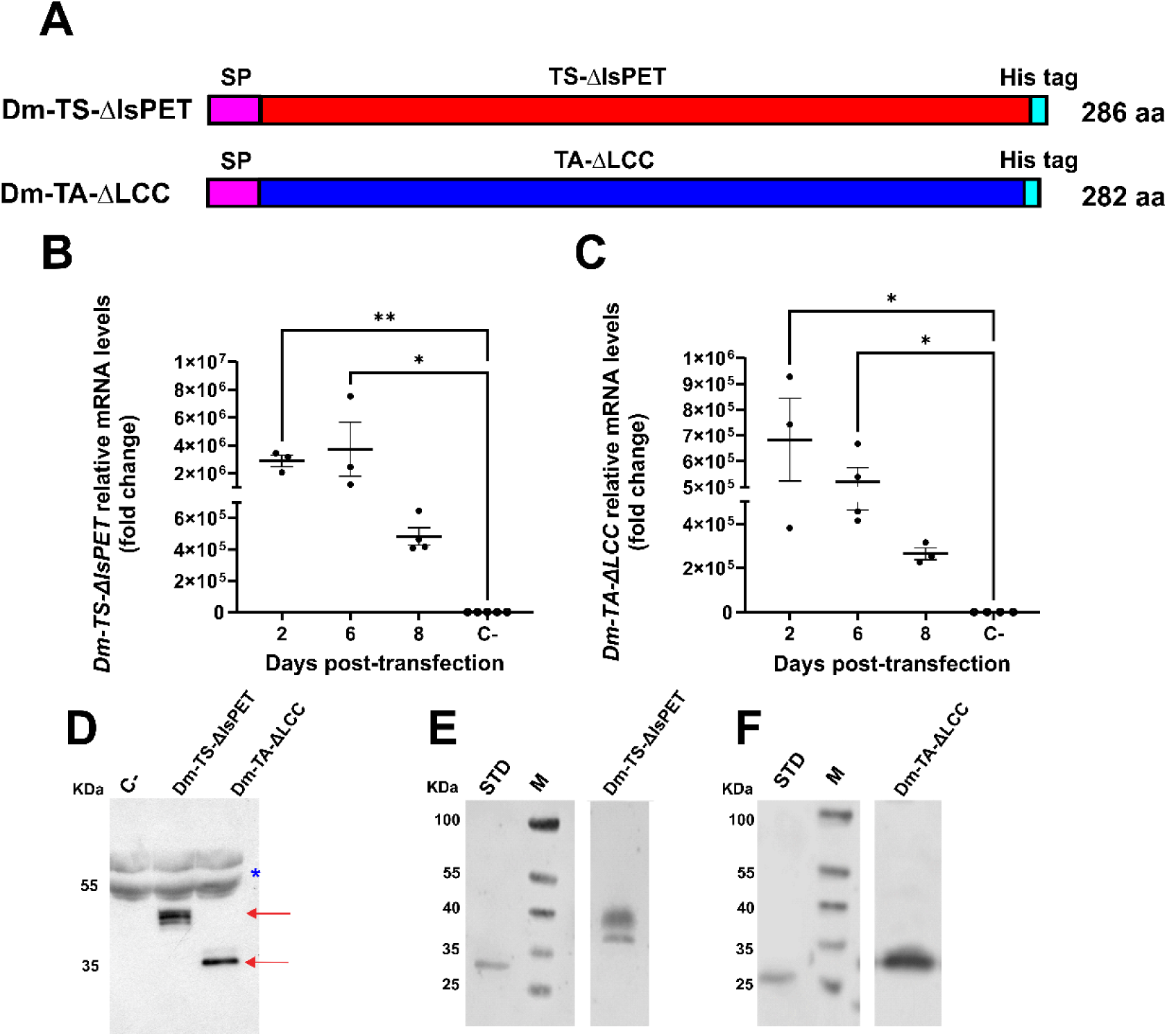
Expression of Dm-TS-ΔIsPET and Dm-TA-ΔLCC in *D. melanogaster* S2R+ cell lines. **A** Schematic representation of the Dm-TS-ΔIsPET and Dm-TA-ΔLCC recombinant proteins. The *in vitro*-evolved 263-residue TS-ΔIsPET and 259-residue TA-ΔLCC variants are shown in red and blue, respectively; the 17-residue Signal Peptide (SP) of a *D. melanogaster* chymotrypsin-like protein (FlyBase ID: FBpp0077758) and the 6-residue His tag (His tag) are shown in magenta and light blue, respectively. aa: amino acids. **B, C** Relative mRNA expression levels (mean ± SEM) of *Dm-TS-ΔIsPET* (**B**) and *Dm-TA-ΔLCC* (**C**) in *Dm-TS-ΔIsPET-* (**B**) and *Dm-TA-ΔLCC-*transfected S2R+ cells (**C**) at 2, 6, and 8 days after transfection. Negative controls (C-) are S2R+ cells transfected with the pAc5-STABLE2-neo vector and sampled at day 2 post-transfection. For each time point, 3-5 biological replicates (dots) were analysed. * and **\*\*** indicate P < 0.05 and P < 0.01, respectively (Kruskal-Wallis test followed by Dunn’s *post hoc* test, comparisons *vs* C-). No statistically significant differences were detected between day 2 and day 6 or 8 (Dunn’s *post hoc* test, comparisons *vs* day 2; P > 0.05). **D-F** Western blots of supernatants from *Dm-TS-ΔIsPET-* (Dm-TS-ΔIsPET) and *Dm-TA-ΔLCC-* (Dm-TA-ΔLCC) transfected S2R+ cells at 8 days post-transfection, developed with anti-His (**D**), anti-TS-ΔIsPET (**E**), and anti-TA-ΔLCC (**F**) antibodies. In (**D**), ∼0.25 mg supernatant proteins per sample have been loaded. C- indicates supernatants from negative controls transfected with the pAc5-STABLE2-neo vector. Red arrows indicate specific anti-His signals detected in Dm-TS-ΔIsPET and Dm-TA-ΔLCC samples, but not in C-. The blue asterisk indicates nonspecific bands visualised with the anti-His antibody in supernatant samples. In (**E**) and (**F**), ∼0.40 mg supernatant proteins per sample were loaded. STD indicates the *E. coli*-produced TS-ΔIsPET (**E**) and TA-LCC (**F**) proteins, used as standards. M: molecular mass marker.

*Dm-TS-ΔIsPET* and *Dm-TA-ΔLCC* mRNAs were efficiently transcribed from the second day after transfection, as revealed by real-time quantitative PCR (qPCR) experiments (Fig. 2B, C). Transcript levels remained detectable at days 6 and 8 and displayed a decrease in mean values over time; however, the difference between day 2 and day 8 was not statistically significant (Fig. 2B, C). Cell viability assessed over a comparable time window showed time-dependent variability across transfected cultures, including empty-vector controls, without statistically significant differences between transfected conditions [two-way repeated measures (RM) ANOVA: P = 0.096, ns; Supplementary Fig. 5]. Accordingly, no construct-specific toxicity was detected under our experimental conditions. Western blot analyses on supernatants of transfected cells using an anti-His antibody showed clear bands corresponding to Dm-TS-ΔIsPET and Dm-TA-ΔLCC at ∼40 and 35 kDa, respectively (Fig. 2D). Using custom monospecific antibodies, a doublet of bands around and below 40 kDa was observed for Dm-TS-ΔIsPET (Fig. 2E) and a band slightly below 35 kDa for Dm-TA-ΔLCC (Fig. 2F). These bands showed a higher molecular mass (M_r_) compared to the *Escherichia coli*-produced TS-ΔIsPET (expected M_r_: 28.8 kDa) and TA-ΔLCC (expected M_r_: 29.0 kDa) variants. These data suggest that both chimeric enzymes could be subjected to post-translational modifications (PTMs) when expressed in the heterologous *D. melanogaster* cell system. One of the most probable PTM is glycosylation, which is known to occur in both cell cultures and whole organism of this insect [47,48]. Consistently, host-dependent modifications such as glycosylation have recently been shown to influence the performance of heterologously expressed wild-type PETase purified from *D. melanogaster* S2 cells [43]. Importantly, both Dm-TS-ΔIsPET and Dm-TA-ΔLCC engineered variants analysed here were actively secreted when expressed in *D. melanogaster* S2R+ cells. Moreover, culture supernatants collected 6 days post-transfection retained activity on p-nitrophenyl acetate (pNPA), a widely used PET esterases model substrate (Supplementary Fig. 6) [49].

### 2.3. Expression of *in vitro-*evolved PET-degrading enzymes in the whole *D. melanogaster* organism

The effects of Dm-TS-ΔIsPET and Dm-TA-ΔLCC expression at the organismal level were evaluated using the Gal4/UAS system [50]. We employed a *daughterless-Gal4* (*daG4*) driver, ubiquitously active from the early developmental stages [51,52], and two *PhiC31* integrase-mediated *Dm-TS-ΔIsPET-*O and *Dm-TA-ΔLCC-*O transgenic lines, carrying a UAS construct for the ectopic expression of Dm-TS-ΔIsPET and Dm-TA-ΔLCC, respectively.

The impact of enzyme expression on post-embryonic development was assessed at 23 and 28 °C by analysing the F1 progeny obtained from mating heterozygous *Dm-TS-ΔIsPET-O*/*CyO* and *Dm-TA-ΔLCC-O*/*TM3* flies to homozygous *daG4* individuals. F1 embryos were monitored until adulthood, when individuals were classified into females and males carrying both the *daG4* driver and UAS construct (*daG4*>*Dm-TS-ΔIsPET-O* and *daG4*>*Dm-TA-ΔLCC-O*), and those carrying the *daG4* driver and the specific balancer chromosome (*daG4/CyO* and *daG4/TM3*). At both temperatures, the number of newly emerged adult flies overexpressing the PET-degrading enzymes (*daG4*>*Dm-TS-ΔIsPET-O* and *daG4*>*Dm-TA-ΔLCC-O*) was comparable to that of their respective controls (*daG4/CyO* and *daG4/TM3*), which did not produce any recombinant enzymes because they lacked the *Dm-TS-ΔIsPET* or *Dm-TA-ΔLCC* transgenes [Fig. 3A and Supplementary Fig. 7A; Fisher’s exact test: P > 0.05, not significant (ns) for all comparisons].

**Fig. 3.**
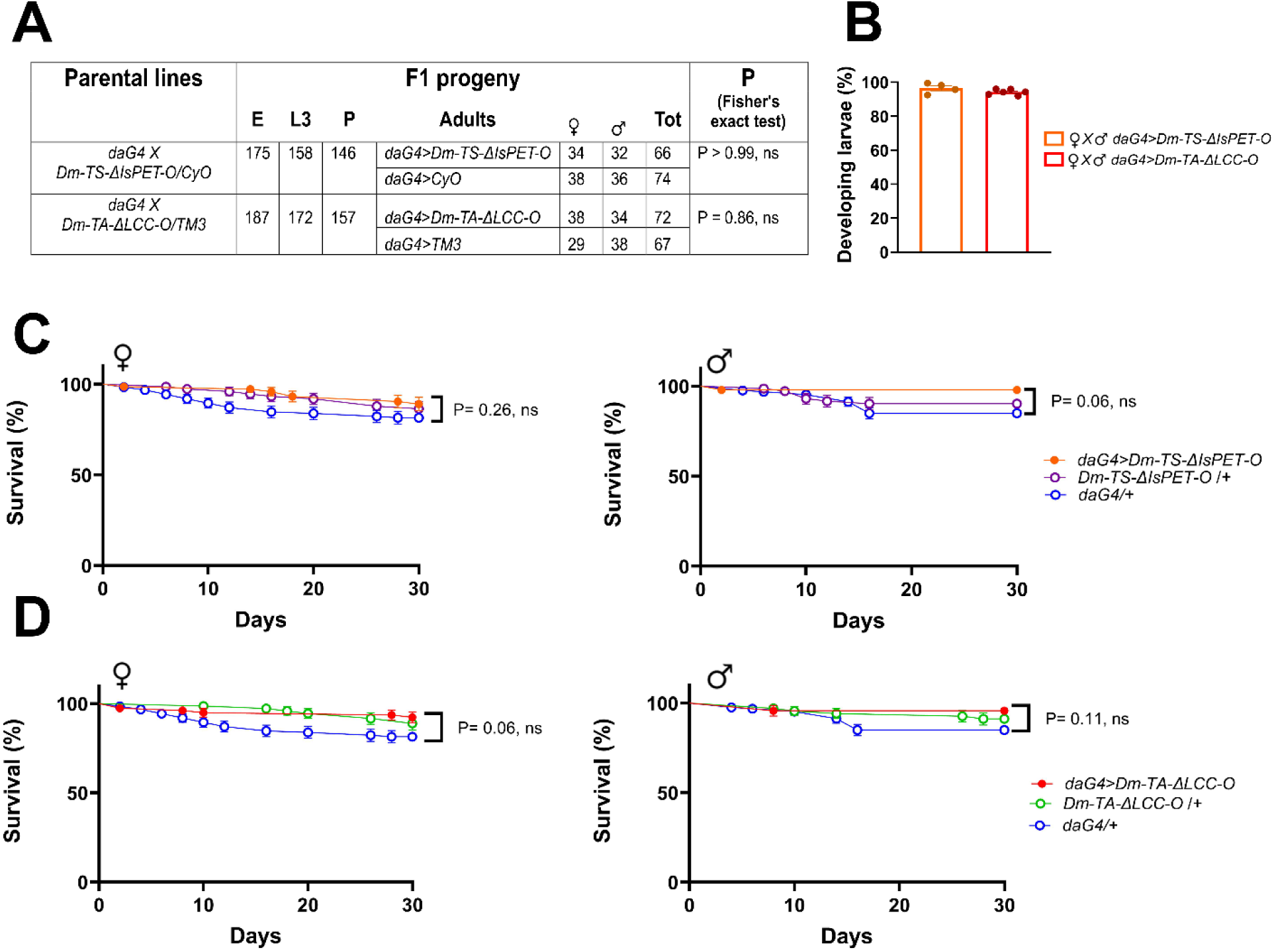
Effects of ubiquitous expression of *Dm-TS-ΔIsPET* and *Dm-TA-ΔLCC* on post-embryonic development, adult fertility, and survival at 23 °C. **A** F1 progeny obtained by mating homozygous *daG4* with heterozygous *Dm-TS-ΔIsPET-O*/*CyO* flies and by mating homozygous *daG4* with heterozygous *Dm-TA-ΔLCC-O*/*TM3* flies. For each cross, total numbers of embryos (E), 3^rd^ instar larvae (L3), and pupae (P) are reported. F1 adult females and males are reported based on their genotype. No significant differences were detected in Fisher’s exact test between female and male *daG4*>*Dm-TS-ΔIsPET-O* flies and female and male *daG4/CyO* controls (P > 0.99, not significant, ns), as well as between female and male *daG4*>*Dm-TA-ΔLCC-O* flies and female and male *daG4/TM3* controls (P= 0.30, ns). **B** Percentage (mean ± SEM) of developing larvae from embryos laid by *daG4*>*Dm-TS-ΔIsPET-O* (orange) and *daG4*>*Dm-TA-ΔLCC-O* (red) flies. Number of analysed embryos: *daG4*>*Dm-TS-ΔIsPET-O*: 649 [4 replicates (dots)]; *daG4*>*Dm-TA-ΔLCC-O*: 771 [6 replicates (dots)]. **C, D** Survival curves (mean % ± SEM) of *daG4*>*Dm-TS-ΔIsPET-O* (**C**) and *daG4*>*Dm-TA-ΔLCC-O* (**D**) flies, compared to their respective negative controls (*daG4/+*, and *Dm-TS-ΔIsPET-O/+* or *Dm-TA-ΔLCC-O/+*). No significant differences in Mantel-Cox test were detected among *daG4*> *Dm-TS-ΔIsPET-O, daG4/+*, and *Dm-TS-ΔIsPET-O/+* flies in both females (left) (P= 0.26, ns) and males (right) (P= 0.06, ns) in (**C**), and among *daG4*>*Dm-TA-ΔLCC-O, daG4/+*, and *Dm-TA-ΔLCC-O/+* flies in both females (left) (P= 0.06, ns) and males (right) (P= 0.11, ns) in (**D**). Number of analysed females and males, respectively: *daG4*>*Dm-TS-ΔIsPET-O:* 73, 47; *daG4/+:* 124, 126; *Dm-TS-ΔIsPET-O/+:* 74, 72; *daG4*> *Dm-TA-ΔLCC-O:* 78, 47; *Dm-TA-ΔLCC-O/+:* 72, 68.

The *daG4*>*Dm-TS-ΔIsPET-O* and *daG4*>*Dm-TA-ΔLCC-O* flies were fertile, with ∼93-95 % of the laid embryos developing into 3^rd^ instar larvae at both 23 and 28 °C (Fig. 3B and Supplementary Fig. 7B). These data indicate that *daG4*-driven activation of either *Dm-TS-ΔIsPET* or *Dm-TA-ΔLCC* transgenes did not compromise gonad functionality and had no negative maternal effects on the development of fertilised eggs.

Comparison of adult survival rates of *daG4*>*Dm-TS-ΔIsPET-O* and *daG4*>*Dm-TA-ΔLCC-O* flies with their respective negative controls, carrying in hemizygosity the *Dm-TS-ΔIsPET* and *Dm-TA-ΔLCC* transgenes (*Dm-TS-ΔIsPET-O/+* and *Dm-TA-ΔLCC-O/*+), and the *daG4* driver (*daG4/+*), revealed no significant variations in lifespan for both sexes, at both temperatures (Fig 3C, D and Supplementary Fig. 7C, D; Log rank test P > 0.05, ns in all comparisons).

We then analysed the expression of the two recombinant PET-degrading enzymes in whole larvae, the developmental stage characterised by intense feeding activity and therefore considered the most suitable stage for insect-mediated PET bioconversion [16,42]. *Dm-TS-ΔIsPET* and *Dm-TA-ΔLCC* chimeric genes resulted highly expressed in *daG4*>*Dm-TS-ΔIsPET-O* and *daG4*>*Dm-TA-ΔLCC-O* 3^rd^ instar larvae in comparison to the negative controls (*daG4/+, Dm-TS-ΔIsPET-O/+,* and *Dm-TA-ΔLCC-O/*+), as revealed by qPCR experiments (Fig. 4A, B). Western blot analyses, using specific anti-TS-ΔIsPET and anti-TA-ΔLCC antibodies, detected Dm-TS-ΔIsPET and Dm-TA-ΔLCC variants as soluble proteins in supernatants derived from *daG4*>*Dm-TS-ΔIsPET-O* and *daG4*>*Dm-TA-ΔLCC-O* larval extracts, respectively (Fig. 4C, D). As already observed in S2R+ cells (Fig. 2E, F), the molecular mass of the larval-derived Dm-TS-ΔIsPET and Dm-TA-ΔLCC proteins was higher than that of the corresponding TS-ΔIsPET and TA-ΔLCC standards, produced in *E. coli* (Fig. 4C, D). In addition, the Dm-TS-ΔIsPET signal showed an evident smear in the *daG4*>*Dm-TS-ΔIsPET-O* larval sample (Fig. 4C). Among the frequent post-translational modifications of ectopically expressed recombinant proteins in *D. melanogaster*, glycosylation results in a higher-than-expected molecular mass of the protein of interest as well as a smear in Western blot analyses. Indeed, computational analysis using the Prosite server (https://prosite.expasy.org/) predicted 8 and 3 potential N-glycosylation sites on TS-ΔIsPET and TA-ΔLCC, respectively [53] (Fig. 4E, F). Based on the most frequent glycosylation patterns in insects’ glycoproteins [54], the resulting predicted molecular masses ranged from 40 to 48 kDa for TS-ΔIsPET and from 31 to 34 kDa for TA-ΔLCC, in agreement with Western blot results [55,56], suggesting that both recombinant variants undergo glycosylation in the larval system.

**Fig. 4.**
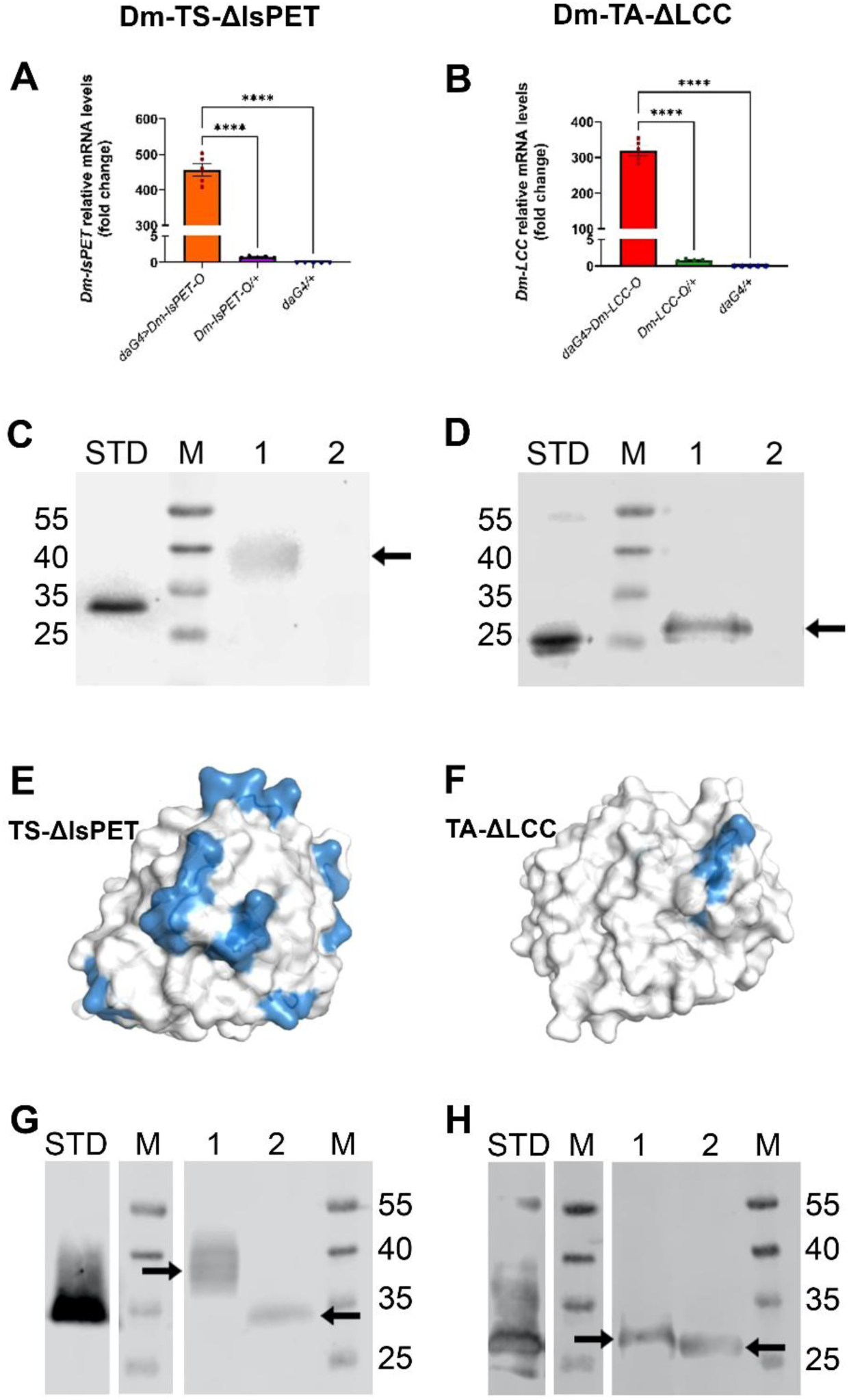
Identification of recombinant Dm-TS-ΔIsPET (left) and Dm-TA-ΔLCC (right) in transgenic larvae. **A, B** mRNA expression of PET hydrolysing enzymes in whole 3^rd^ instar larvae. Relative mRNA levels (mean ± SEM) of *Dm-TS-ΔIsPET* in *daG4*>*Dm-TS-ΔIsPET-O* and negative controls (*daG4/+* and *Dm-TS-ΔIsPET-O/+*) (**A**), and *Dm-TA-ΔLCC* in *daG4*>*Dm-TA-ΔLCC-O* and negative controls (*daG4/+* and *Dm-TA-ΔLCC-O/+*) (**B**). Five biological replicates (dots), each with 10 larvae, per genotype were analysed. **** indicates P < 0.0001, significant differences in the relative expression levels determined with Brown-Forsythe ANOVA (F_2,4_ = 656.8) followed by Dunnett’s multiple comparisons test. **C, D** Western blots of larval extracts using anti-TS-ΔIsPET or anti-TA-ΔLCC antibodies. **C** Extracts from *Drosophila* expressing Dm-TS-ΔIsPET. STD: 50 ng of bacterial recombinant TS-ΔIsPET; M: molecular mass standards; 1: ∼0.3 mg of extract proteins from *daG4>Dm-TS-ΔIsPET-O* larvae; 2: ∼0.3 mg of extract proteins from *Dm-TS-ΔIsPET*-*O/+* negative controls. **D** Extracts from *Drosophila* expressing Dm-TA-ΔLCC. STD: 50 ng of bacterial recombinant TA-ΔLCC; M: molecular mass standards; 1: 20 μL extract (∼1.3 mg proteins) from *daG4>Dm-TA-ΔLCC-O* larvae; 2: 20 μL extract (∼1.3 mg proteins) from *Dm-TA-ΔLCC-O/+* negative controls. **E, F** Potential N-glycosylation sites on the surface of recombinant TS-ΔIsPET (**E**) and TA-ΔLCC (**F**) as predicted by the PROSITE server (https://prosite.expasy.org/). Proteins are shown with surface representation. Putative N-glycosylation sites are shown in blue. 3D models of the proteins were produced using I-Tasser [55]. Images were prepared with PyMOL [56]. **G H** N-deglycosylation using PNGase F of Dm-TS-ΔIsPET (**G**) and Dm-TA-ΔLCC (**H**) expressed in *Drosophila*. STD: Recombinant protein expressed in bacteria (0.5 μg); M: molecular mass standards; 1: protein before deglycosylation; 2: protein after deglycosylation. Black arrows indicate proteins of interest.

To experimentally assess glycosylation of the recombinant proteins, supernatants from *daG4*>*Dm-TS-ΔIsPET-O* and *daG4*>*Dm-TA-ΔLCC-O* larval extracts were subjected to *in vitro* enzymatic deglycosylation using PNGase F to remove sugar residues at the N-amino glycosylation sites. Western blot analysis showed a decrease in molecular mass of ∼6 and ∼2 kDa for Dm-TS-ΔIsPET and Dm-TA-ΔLCC, respectively (Fig. 4G, H). Specifically, the original molecular mass of Dm-TS-ΔIsPET (M_r_: 40-45 kDa) and Dm-TA-ΔLCC (M_r_: ∼28 kDa) shifted to 34 and ∼26 kDa, respectively, after PNGase F incubation (Fig. 4G, H), confirming glycosylation of both variants [47], thus providing direct experimental evidence of glycosylation in a whole-organism context.

### 2.4. Enzymatic activities of Dm-TS-ΔIsPET and Dm-TA-ΔLCC in D. *melanogaster* larvae

To test whether Dm-TS-ΔIsPET and Dm-TA-ΔLCC were enzymatically active in *daG4*>*Dm-TS-ΔIsPET-O* and *daG4*>*Dm-TA-ΔLCC-O* larvae, we assessed their activity on pNPA [49]. Larval extracts from negative controls (*daG4/+, Dm-TS-ΔIsPET-O/+,* and *Dm-TA-ΔLCC-O/*+) showed a basal esterase activity ranging from 1.36 ± 0.10 to 2.05 ± 0.16 U/g_larvae_, respectively (Fig. 5). Extracts from *daG4*>*Dm-TS-ΔIsPET-O* (expressing Dm-TS-ΔIsPET) and *daG4*>*Dm-TA-ΔLCC-O* (expressing Dm-TA-ΔLCC) individuals showed an esterase activity of 3.41 ± 0.06 and 11.02 ± 0.94 U/g_larvae_, respectively, corresponding to a net increase of 1.58 ± 0.05 and 8.97 ± 0.32 U/g_larvae_, compared to the basal activity of the respective *Dm-TS-ΔIsPET-O/+* and *Dm-TA-ΔLCC-O/*+ negative controls.

**Fig. 5.**
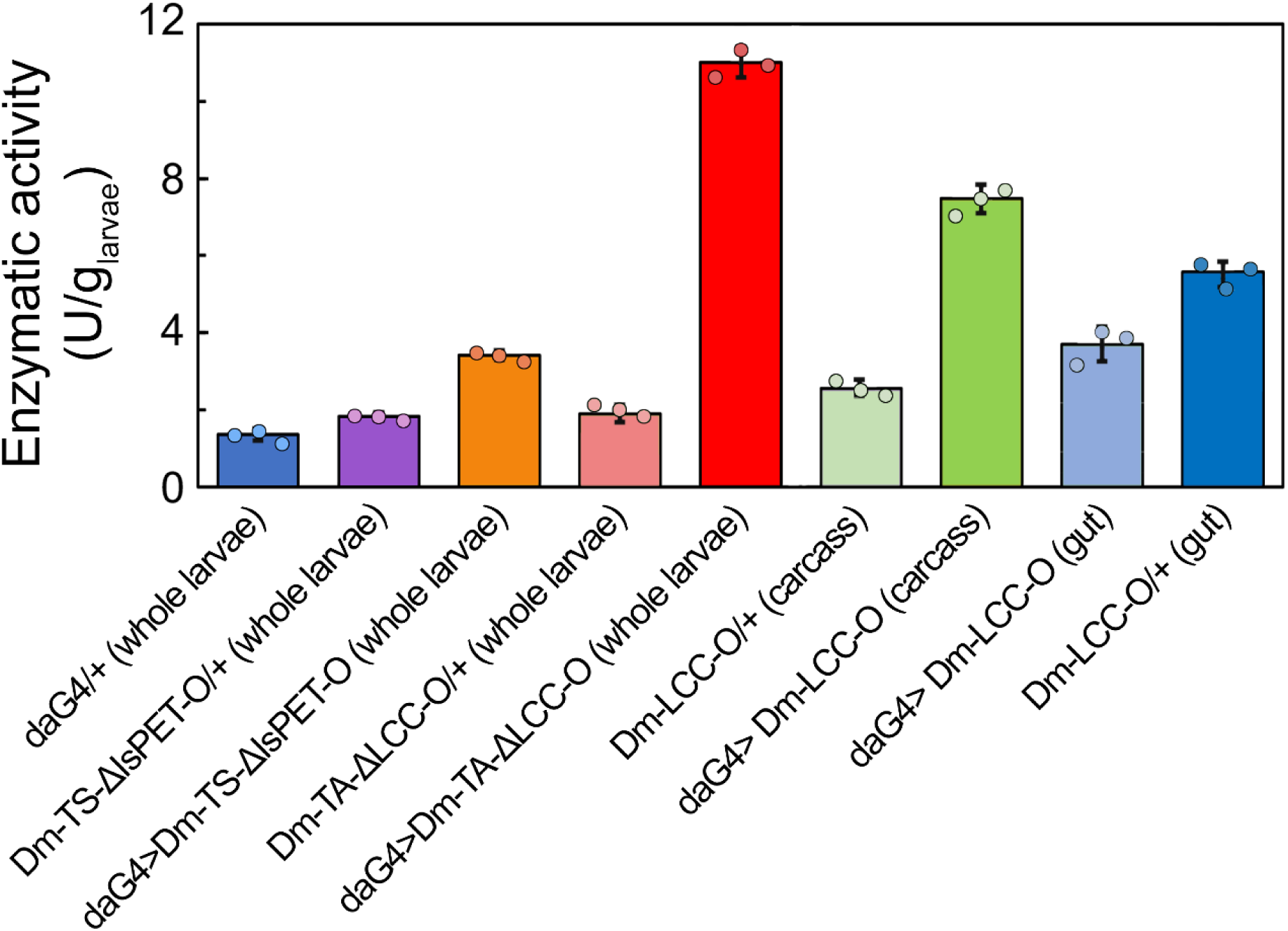
Esterase activity in larval extracts. Enzymatic units of esterases in extracts from *daG4>Dm-TS-ΔIsPET-O* whole larvae (orange), *daG4>Dm-TA-ΔLCC-O* whole larvae (red), *daG4>Dm-TA-ΔLCC-O* carcasses (pale red) and *daG4>Dm-TA-ΔLCC-O* guts (blue). Corresponding negative controls (not expressing the recombinant enzymes): *daG4/+* whole larvae (grey), *Dm-TS-ΔIsPET-O/+* whole larvae (purple), *Dm-TA-ΔLCC-O/+* whole larvae (green), *daG4>Dm-TA-ΔLCC-O* carcasses (pale green) and *daG4>Dm-TA-ΔLCC-O* guts (pale blue). For each genotype, the mean ± SEM of 3 replicates (dots) is shown.

These data indicate that the increased esterase activity was due to the heterologous expression of the two recombinant PET hydrolases in larval tissues (Fig. 5).

The estimated amounts of insect-produced recombinant Dm-TS-ΔIsPET and Dm-TA-ΔLCC were 238 ± 26 and 138 ± 40 μg/g_larvae_, respectively (estimated from the variants’ specific activity, Supplementary Fig. 8). Since the net esterase activity of *Drosophila-*expressed Dm-TA-ΔLCC was almost than 6-fold higher than that of Dm-TS-ΔIsPET, the *daG4*>*Dm-TA-ΔLCC-O* larval extracts were used for further investigations. To assess whether the esterase activity of Dm-TA-ΔLCC was also specifically detectable in larval guts, these were isolated and analysed separately from the rest of the body (hereafter referred to as carcasses), comprising epidermis, brain, salivary glands, fat body, Malpighian tubules, and tracheae.

The presence of the recombinant protein in gut extracts from *daG4*>*Dm-TA-ΔLCC-O* larvae was detected by Western blot analysis (Supplementary Fig. 9). In the same samples, esterase activity was observed (5.53 U/g_gut_; Fig. 5), demonstrating enzymatic activity in the gut. The activity measured in the remaining carcass tissues (7.44 U/g_carcass_) reflects the combined contribution of non-intestinal tissues expressing the transgene under the control of the ubiquitous *daG4* driver. In control larvae, the mean basal esterase activity was slightly higher in gut extracts than in the corresponding carcass tissues (3.69 U/g_gut_ vs 2.57 U/g_carcass_, respectively), likely reflecting the contribution of endogenous gut digestive enzymes (Fig. 5).

The performance of the Dm-TA-ΔLCC variant on amorphous PET nanoparticles’ degradation was evaluated incubating ∼240 μg PET nanoparticles per biodegradation reaction at 55 °C, a temperature below the glass transition temperature (T*_g_*) of PET, in the presence of larval extracts. Degradation reactions were performed at pH 9, as the enzyme exhibits greater activity under basic conditions [57]. The depolymerisation of PET was determined by measuring the accumulation of soluble degradation products (i.e., MHET and TPA) in the reaction mixture. Analysis of absorption spectra showed a time-dependent increase of a peak centered at ∼246 nm corresponding to the absorbance peak of MHET and TPA [49] (Fig. 6 and Supplementary Fig. 10). The amount of soluble depolymerisation products at each time point was determined by HPLC analysis. Chromatograms showed two main peaks at an elution time of 9.6 and 10.8 min corresponding to TPA and MHET, respectively. The relative concentration of the two depolymerisation products changed during conversion: TPA represented ∼33 % of the total amount of products at the beginning of the reaction and ∼96 % at the end (i.e., after 6 and 144 h, respectively) (Fig. 6A). This is due to the fact that LCC catalysed two sequential reactions: the hydrolysis of PET to MHET (faster reaction) and the hydrolysis of MHET to TPA (slower reaction). The highest concentration of depolymerisation products was 118 ± 24 μM, obtained after 144 h of incubation (Fig. 6B), which corresponds to the depolymerisation of ∼10.3 % of the initial amount of PET nanoparticles. The highest rate of PET hydrolysis was 66.6 μmol_products_/g_extract proteins_/day (measured after 3 h of reaction). This corresponds to about 13.5 μmol_products_/day of degraded PET per 1 g of Dm-TA-ΔLCC*-*overexpressing larvae. Moreover, gut extracts from *Dm-TA-ΔLCC*-overexpressing larvae were able to depolymerise PET nanoparticles. Under these conditions, the highest concentration of depolymerisation products was 79.3 ± 7.3 μM, obtained after 144 h of incubation (Fig. 6B), which corresponds to the depolymerisation of ∼6.9 % of the initial amount of PET nanoparticles. In this case, the highest rate of PET hydrolysis was 125.6 μmol_products_/g_extract proteins_/day (measured after 3 h of reaction), corresponding to ∼13.6 μmol_products_/day of degraded PET per 1 g of guts from *Dm-TA-ΔLCC-*overexpressing larvae.

**Fig. 6.**
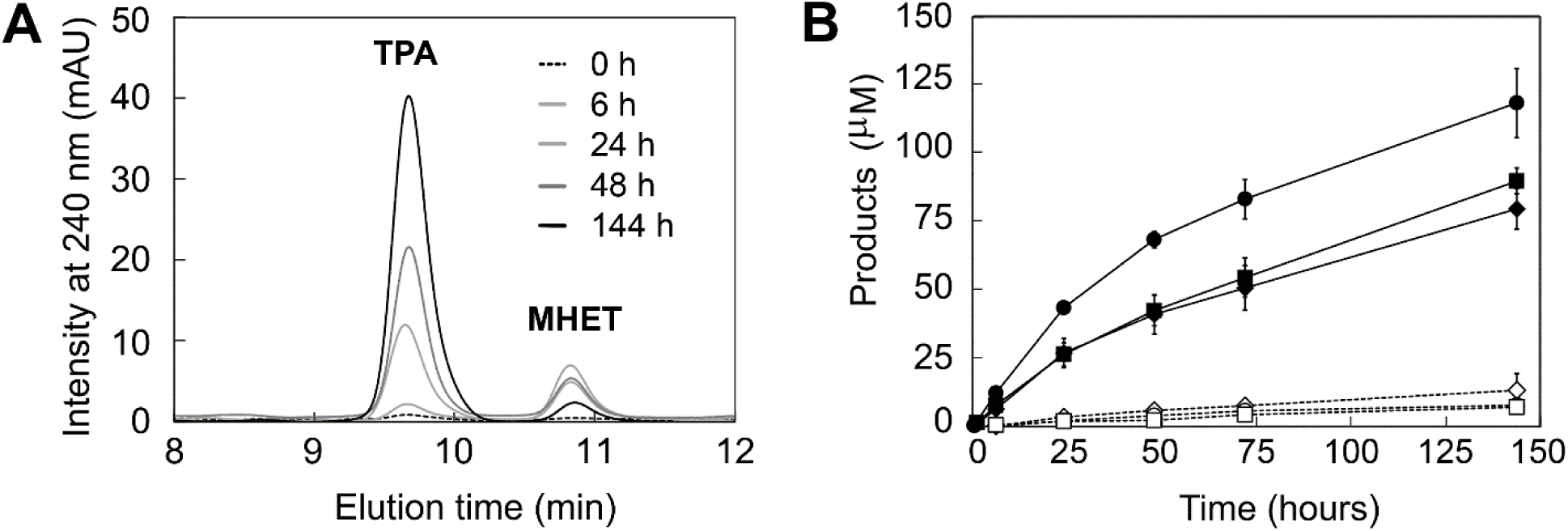
Depolymerisation of PET nanoparticles by extracts from larvae expressing Dm-TA-ΔLCC. **A** HPLC chromatograms of the reaction mixture at increasing reaction times. **B** Kinetics of accumulation of soluble PET depolymerisation products during bioconversion catalysed by different extracts from *Dm-TA-ΔLCC*-expressing larvae (*daG4*>*Dm-TA-ΔLCC-O*) (closed symbols) and corresponding negative controls (*Dm-TA-ΔLCC-O/*+) (open symbols). Larval, gut, and carcass extracts are shown as circles, diamonds, and squares, respectively. In **B** the mean ± SEM of 3 replicates are shown.

### 2.5. Tissue-specific expression and functional impact of Dm-TA-ΔLCC in the larval gut

Given its ability to hydrolyse PET, Dm-TA-ΔLCC was further analysed at the intestinal level. Transcription of *Dm-TA-ΔLCC* mRNA in the guts of *daG4>Dm-TA-ΔLCC-O* larvae was confirmed by qPCR (Supplementary Fig. 11). Subsequently, we investigated the expression pattern of Dm-TA-ΔLCC protein via immunolocalisation analyses (Fig 7). The presence of the Dm-TA-ΔLCC recombinant enzyme throughout the length of the alimentary canal in *daG4*>*Dm-TA-ΔLCC-O* larvae was demonstrated by whole-mount staining (Fig. 7G-H), as well as through immunogold labelling (Fig. 7I). Both results also proved Dm-TA-ΔLCC localisation at both epithelial and luminal level compared to controls (*daG4/+* and *Dm-TA-ΔLCC-O/+*) (Fig. 7A-F).

**Fig. 7.**
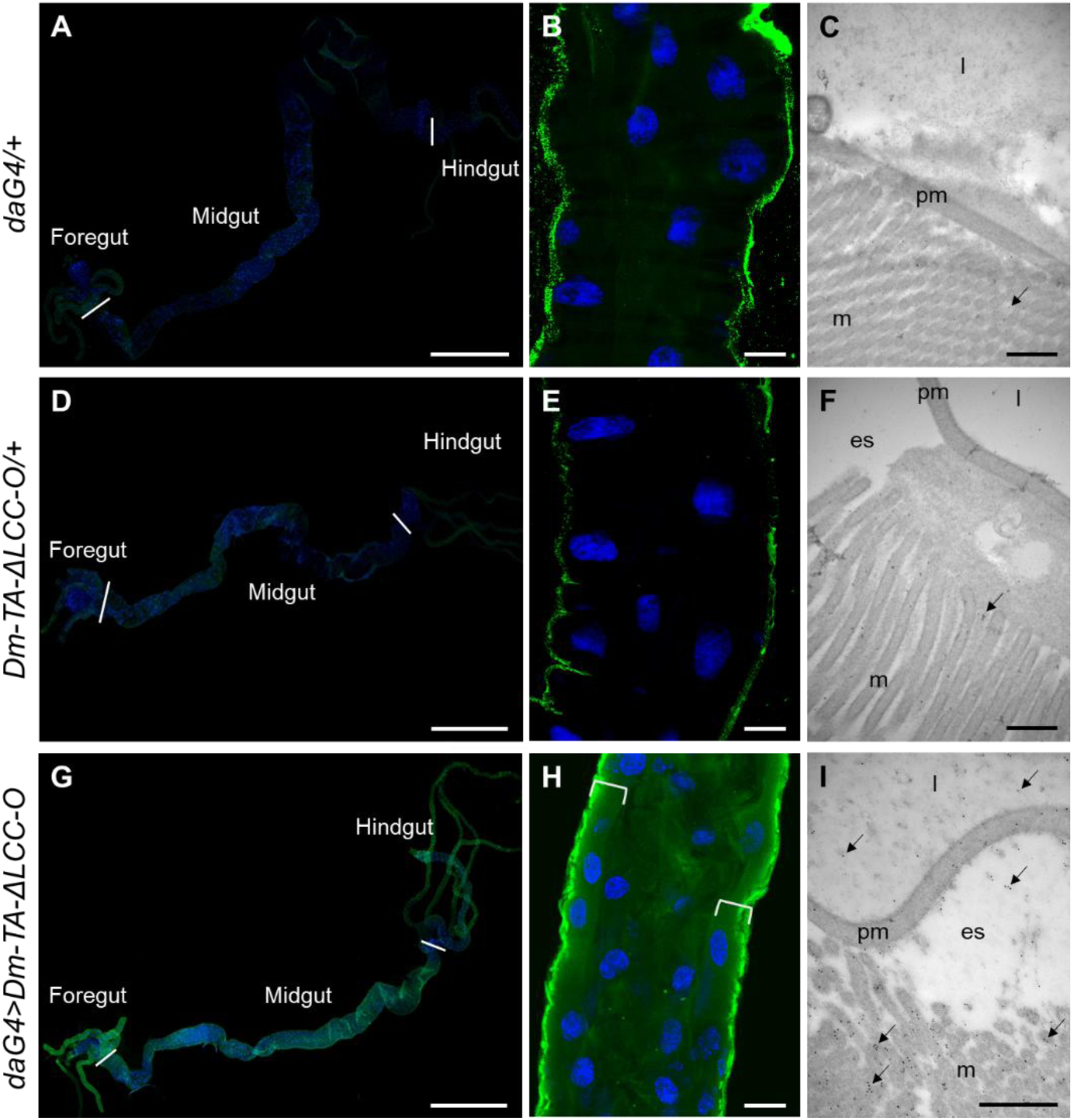
Expression of the Dm-TA-ΔLCC recombinant enzyme in the whole gut of *daG4*>*Dm-TA-ΔLCC-O* larvae. Localisation (green signal) of Dm-TA-ΔLCC by whole-mount staining (**A-B, D-, E, G-H**) and immunogold labelling on cross-sections (**C, F, I**) using gut samples isolated from *daG4/+* (**A-C**), *Dm-TA-ΔLCC-O/+* (**D-F**), and *daG4*>*Dm-TA-ΔLCC-O* (**G-I**) larvae. Brackets: midgut epithelium; es: ectoperitrophic space; l: lumen; m: microvilli; pm: peritrophic matrix. Nuclei are stained in blue (DAPI). Arrows indicate gold particles (**C, F, I**). Bars: 1 mm (**A, D, G**), 20 μm (**B, E, H**), 500 nm (**C, F, I**).

To determine whether Dm-TA-ΔLCC production affected the larval gut morphology, ultrastructural analysis of the *Dm-TA-ΔLCC*-overexpressing midguts was performed via Transmission Electron Microscopy (TEM) analysis. The morphology of the midgut epithelium in *daG4*>*Dm-TA-ΔLCC-O* larvae (Fig. 8C) was comparable to that of negative controls (*Dm-TA-ΔLCC/+* and *daG4/+*) (Fig. 8A, B). No ultrastructural modifications were identified in the apical membrane of gut cells in *Dm-TA-ΔLCC*-overexpressing larvae, since a well-developed brush border with microvilli was recognisable in the epithelium of both *daG4*>*Dm-TA-ΔLCC-O* larvae (Fig. 8F) and negative controls (Fig. 8D, E). Similarly, the basal region of the gut cells showed a well-organised ultrastructure as confirmed by a continuous basal lamina, as well as muscle cells in both *Dm-TA-ΔLCC*-overexpressing larvae (Fig. 8I) and controls (Fig. 8G, H).

**Fig. 8.**
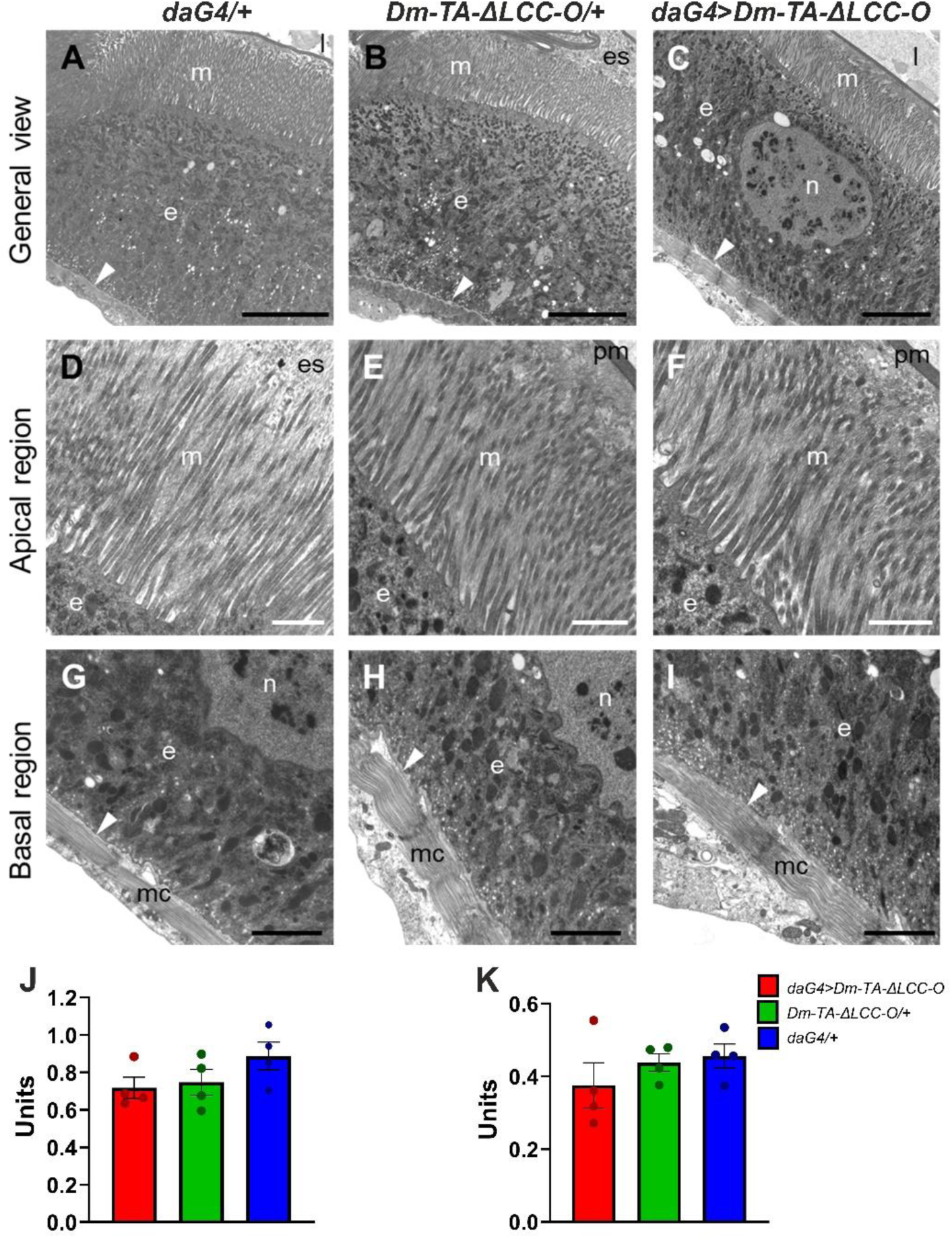
Effects of ectopic *Dm-TA-ΔLCC* expression on midgut ultrastructure and digestive activity in 3^rd^ instar larvae. **A-I** TEM analysis of midgut cells in *daG4/+* (**A, D, G**), *Dm-TA-ΔLCC-O/+* (**B, E, H**) controls and *daG4*>*Dm-TA-ΔLCC-O* larvae (**C, F, I**). Arrowhead: basal lamina; e: epithelium; es: ectoperitrophic space; l: lumen; m: microvilli; mc: muscle cells; n: nucleus; pm: peritrophic matrix. Bars: 5 μm (**A-C**), 2 μm (**G-I**), 1 μm (**D-F**). **J, K** Enzymatic activities (mean ± SEM) of APN (**J**) and ALP (**K**) in homogenates of the midgut from *daG4*>*Dm-TA-ΔLCC-O* larvae and negative controls (*Dm-TA-ΔLCC-O/+* and *daG4/+)*. For each genotype, four biological replicates (dots), each with at least three technical replicates, were analysed. Both enzymatic activities were not statistically different among genotypes [one-way ANOVA: F_2,9_ = 1.84; P = 0.21, ns (**J**); F_2,9_ = 0.96; P = 0.42, ns (**K**)].

Potential effects of a *Dm-TA-ΔLCC* expression on larval midgut function were evaluated by measuring the activity of aminopeptidase N (EC 3.4.11.2) (APN) and alkaline phosphatase (EC 3.1.3.1) (ALP), two enzymes associated with the brush border membranes of insect midgut cells [58]. APN and ALP are well-established marker enzymes of microvilli of digestive absorbing epithelia in insects of different orders, including Diptera [58–61]. APN and ALP activities measured in midgut homogenates from *daG4*>*Dm-TA-ΔLCC-O* larvae were not statistically different from those of both *Dm-TA-ΔLCC-O/+* and *daG4/+* negative controls (Fig. 8J, K), suggesting that the production of the TA-ΔLCC does not affect midgut digestive functions in *D. melanogaster* larvae.

### 2.6. Effects of PET nanoparticle exposure in *Dm-TA-ΔLCC* expressing larvae

Lastly, we evaluated whether exposure to PET nanoparticles affects fitness parameters in larvae expressing the Dm-TA-ΔLCC variant. To this end, we compared post-embryonic development of *daG4*>*Dm-TA-ΔLCC-O* individuals and the corresponding negative controls (*Dm-TA-ΔLCC-O/*+; *daG4/+*) when reared at 28 °C on food supplemented with 0.05% PET nanoparticles and on respective PET-free control food (Fig 9). This concentration corresponds to the highest dose previously tested in *D. melanogaster* at 25 °C, in which wild-type larvae were shown to ingest and internalise PET nanoparticles without evident effects on egg-to-adult viability [62], and was applied here at 28 °C as a more stringent growth condition.

**Fig. 9.**
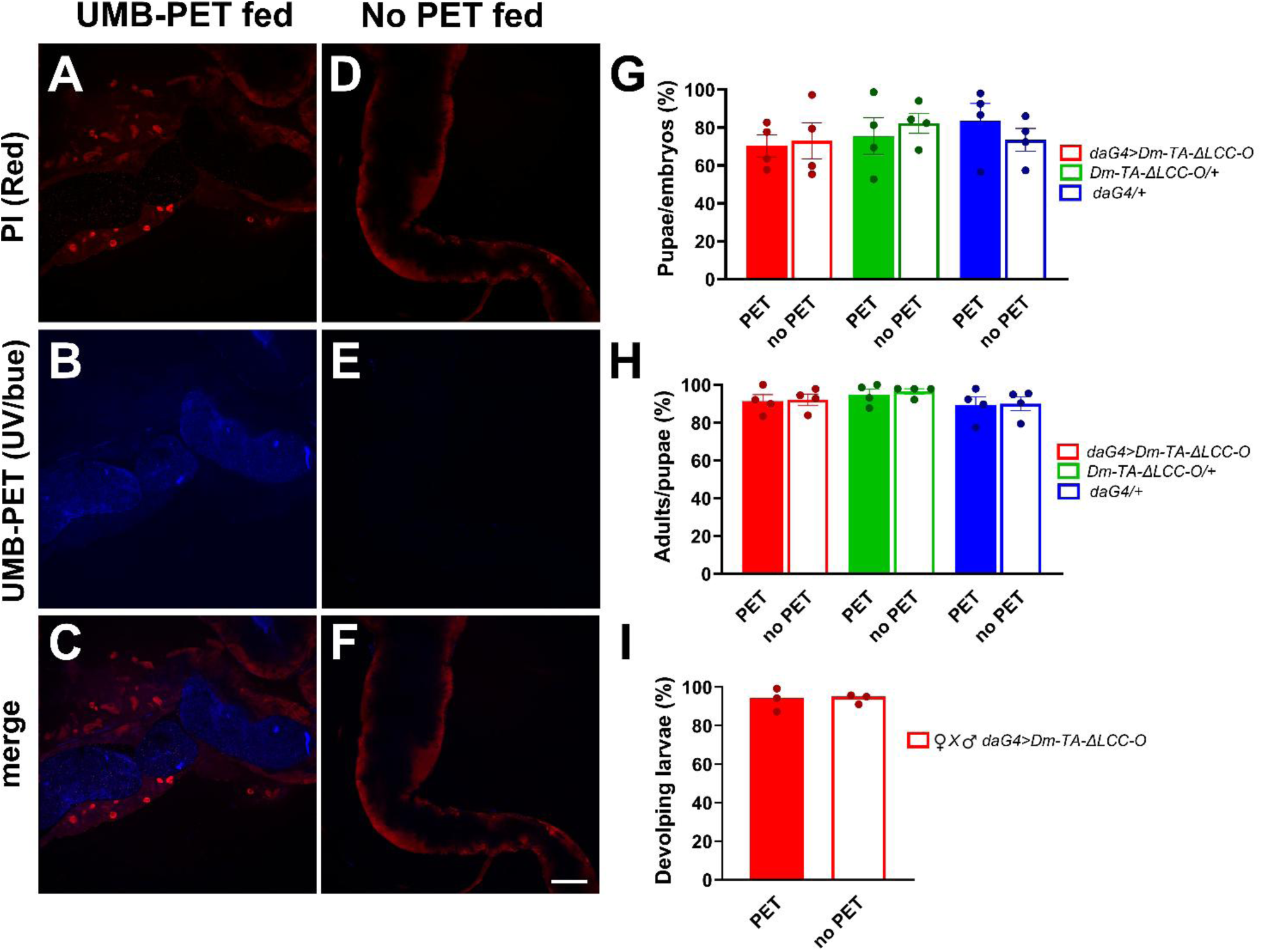
Effects of PET nanoparticle exposure on post-embryonic development and adult fertility in *D. melanogaster* overexpressing *Dm-TA-ΔLCC* at 28 °C. Visualisation of PET nanoparticle ingestion in the larval midgut of *D. melanogaster* reared on a diet supplemented with 0.05 % umbelliferone-stained PET nanoparticles (UMB-PET fed) (**A-C**) and on control diet without PET nanoparticles (No PET fed) (**D-F**). Guts were stained with propidium iodide (PI, red) (**A, D**) and imaged for UMB-PET fluorescence (blue) (**B, E**); merged images are shown in (**C, F**). Bar: 100 μm (reported in **F**). **G**, **H** Percentage (mean ± SEM) of pupae developed from embryos (**G**) and adults emerged from pupae (**H**) belonging to the *daG4>Dm-TA-ΔLCC-O* (red), *Dm-TA-ΔLCC-O/+* (green) and *daG4/+* (blue) genotypes, reared on a diet supplemented with 0.05% PET nanoparticles (filled bars) and on a control diet without PET nanoparticles (open bars). For each genotype and condition, 4 biological replicates (dots) were analysed, each including 45-80 embryos (**G**) or 26-71 pupae (**H**). No statistically significant differences were observed among genotypes or dietary conditions [two-way ANOVA: F₂,₁₈ = 0.76; P = 0.48, ns (**G)**; F₂,₁₈ = 0.012; P= 0.98, ns (**H**)]. **I** Percentage (mean ± SEM) of developing larvae from embryos laid by adults belonging to the *daG4*>*Dm-TA-ΔLCC-O* genotype reared on a diet supplemented with 0.05% PET nanoparticles (filled bars) and on a control diet without PET nanoparticles (open bars). For each condition, 3 biological replicates (dots) were analysed, each including 67-170 embryos. No statistically significant difference was observed between conditions (unpaired two-tailed t-test: P = 0.91, ns).

Larval ingestion of PET nanoparticles was verified using PET nanoparticles labelled with the blue fluorophore umbelliferone (UMB-PET). After chronic exposure from the embryonic stage to the 3^rd^ larval instar, a blue fluorescence signal was detected in the midgut lumen of 0.05% UMB-PET-fed larvae, whereas only weak background fluorescence was observed in the corresponding controls reared on a PET-free diet (Fig. 9 A–F).

We then analysed post-embryonic developmental performance by determining the percentage of pupae developing from 24-h-old embryos and of adults emerging from pupae in *daG4>Dm-TA-ΔLCC-O* individuals and the respective negative controls (*Dm-TA-ΔLCC-O/*+; *daG4/+*), reared on PET-supplemented or PET-free control food (Fig. 9G, H). For all analysed genotypes, no significant differences were detected between PET-fed and control groups, with no significant genotype × diet interaction (two-way ANOVA: P = 0.48 and 0.98 for Fig. 9G and H, respectively).

Finally, to assess whether chronic exposure to PET nanoparticles during development might affect fertility, *daG4*>*Dm-TA-ΔLCC-O* males and females that had developed in the presence or absence of PET nanoparticles were crossed, and progeny viability was quantified (Fig. 9I). The percentage of developing larvae from embryos laid by PET-exposed adults was not statistically different from that of controls, indicating that developmental exposure to PET nanoparticles did not impair adult fertility in *Dm-TA-ΔLCC* overexpressing flies (unpaired two-tailed t-test: P = 0.91; Fig 9I).

## 3. Discussion

Organic feedstocks, such as OFMSW, are often contaminated with micro- and nanoplastics, including PET [63,64]. Insect larvae have emerged as powerful bioreactors to convert organic waste into high-quality biomolecules [15–18]. Although some insects can ingest plastic polymers and show partial degradation, these processes remain variable and context-dependent [24,28]. Conversely, engineered enzymes of bacterial origin can efficiently hydrolyse PET into its monomers [32], suggesting the potential of combining insect-based systems with heterologous expression of plastic-degrading enzymes.

To assess whether insects tolerate the functional expression of *in vitro-*evolved PET-degrading enzymes, we generated transgenic *D. melanogaster* lines early and ubiquitously expressing Dm-TS-ΔIsPET and Dm-TA-ΔLCC. These lines showed normal development and fertility, with no detectable deleterious phenotypes. Ultrastructural and biochemical analyses confirmed that Dm-TA-ΔLCC synthesis did not alter midgut morphology and digestive activity. Together, these results indicate that *D. melanogaster* can sustain the expression of engineered PET-hydrolysing enzymes. Consistently, a recent independent study reported comparable tolerance following expression of wild-type PETase in transgenic flies [43].

In addition, we demonstrated that the signal peptide of a chymotrypsin-like protein was effective in directing extracellular secretion of Dm-TS-ΔIsPET and Dm-TA-ΔLCC in *D. melanogaster* S2R+ cells and *in vivo*, particularly in the larval gut. This signal peptide thus expands the repertoire of tools available for heterologous protein secretion in *D. melanogaster* [42].

As previously reported for PET-degrading enzymes in yeast expression systems [65–67], Dm-TS-ΔIsPET and Dm-TA-ΔLCC were glycosylated when expressed in *D. melanogaster*. This PTM did not abolish the activity of the insect-expressed enzymes. Dm-TA-ΔLCC showed an almost 6-fold higher activity than Dm-TS-ΔIsPET, establishing it as a substantially more effective biocatalyst in this insect host. This difference likely results from a combination of factors, including different glycosylation effects on enzyme properties, the presence of disulfide bridges in Dm-TS-ΔIsPET but not in Dm-TA-ΔLCC, and/or differential interactions with components of the larval extracts. Indeed, glycosylated yeast-expressed LCC shows improved performance (PET-hydrolysing activity and stability) in comparison to its bacterial counterpart, while the effects of this post-translational modification on IsPETase appear more complex and context-dependent [65–67]. The expression of wild-type PETase in *D. melanogaster* S2 cells has also highlighted the occurrence and functional impact of glycosylation, showing that this modification enhances stability over time, whereas the deglycosylated form displays higher initial activity under short-term *in vitro* conditions using purified enzyme [43].

Based on the comparative activity observed on the model substrate pNPA, crude extracts from whole-body and gut tissues of Dm-TA-ΔLCC-expressing larvae were further evaluated, demonstrating the ability to depolymerise PET nanoparticles when tested *in vitro* under optimal conditions. A direct comparison of PET-depolymerisation efficacy between Dm-TA-ΔLCC and previously reported PET-degrading enzyme variants is not straightforward due to differences in heterologous expression systems (e.g., prokaryotic hosts, yeast, cell cultures such as *Drosophila* S2 cells, or whole transgenic organisms), glycosylation status, formulation (e.g., purified enzyme vs. crude *Drosophila* extract), and assay conditions (e.g., buffer composition, temperature, and PET substrate). In our system, the amount of TA-ΔLCC in *D. melanogaster* larvae (138 ± 40 μg/g_larvae_) was ∼14-fold lower than that obtained in *E. coli* (∼2.0 mg/g_cells_). However, this difference should not be interpreted exclusively in terms of protein yield. Transgenic insects should not be considered as bioreactors for protein production, but as living systems in which enzyme availability is achieved *via* persistent tissue-specific expression and secretion. This highlights organism-level features relevant to plastic-contaminated waste processing, where enzyme activity is integrated with feeding on complex substrates. Interestingly, heterologous expression of wild-type PETase in *D. melanogaster* allows detectable PET hydrolysis *in vivo*, which is slow but continuous and strongly influenced by environmental factors such as pH and substrate accessibility [43].

*D. melanogaster* overexpressing the Dm-TA-ΔLCC variant did not display detectable developmental or reproductive impairments, even when chronically exposed to PET nanoparticles. These data indicate that Dm-TA-ΔLCC production together with PET ingestion is well tolerated even at 0.05% PET nanoparticle concentration, which exceeds environmental levels recently reported for micro- and nanoplastics [7,68].

These observations provide a rationale for exploring the expression of engineered PET-hydrolysing enzymes in insects currently investigated for waste bioconversion [28]. Among them, *H. illucens* represents a promising candidate for its ability to grow on a wide range of organic waste, including OFMSW [15,16,19]. The recent development of both conventional transgenesis and advanced genome editing approaches in *H. illucens* [69-72] provides a basis for translating the approach described here to this species. As both *H. illucens* and *D. melanogaster* belong to the order Diptera and suborder Brachycera, key aspects of the fruit fly experimental paradigm, including efficient gut secretion and host tolerance to PET-degrading enzyme production, may be transferable, although this remains to be experimentally validated. In line with this, PET-degrading enzyme overexpression did not result in evident adverse effects in *D. melanogaster* at both 23 and 28 °C. Furthermore, the higher activity of TA-ΔLCC relative to TS-ΔIsPET in the *Drosophila* host, together with its exclusive production of TPA as the final depolymerisation product, makes it a particularly attractive candidate for translation to *H. illucens*, where complete depolymerisation to monomers is a prerequisite for a potential downstream valorisation.

These observations suggest that *H. illucens* transgenic lines stably inheriting a functional TA-ΔLCC variant may be viable and fertile. *H. illucens* larvae are typically reared at 28-30 °C; however, under mass-rearing conditions, larval aggregation can increase substrate temperature by ∼10 °C, frequently reaching 40-45 °C [73]. This temperature range is closer to the temperature required for an efficient PET depolymerisation by the TA-ΔLCC variant observed *in vitro*. Under this scenario, intestinal-driven synthesis and secretion of TA-ΔLCC may contribute to enzyme availability in the gut lumen during feeding. In this context, *H. illucens* represents a promising host for further development toward applied scenarios. However, assessment of PET hydrolysis efficiency and the fate of degradation products, including potential bioaccumulation and assimilation, will be required. These engineered *H. illucens* lines could also provide a basis for the development of confined large-scale production strains, suitable for mass rearing on PET-contaminated OFMSW and integrated with genetic containment strategies to minimise the potential risks associated with accidental release or ecological spillover during mass rearing [74].

## 4. Conclusions

This study demonstrates that *D. melanogaster* is a tractable system for the *in vivo* expression of engineered PET-degrading enzymes, allowing the comparative evaluation of biochemically distinct variants. Together with recent independent findings on wild-type PETase [43], our results support the use of this model for the exploration of engineered PET-degrading enzymes in an insect context.

To our knowledge, the TA-ΔLCC variant represents the first LCC-derived enzyme functionally evaluated in an insect host. This variant is a more active biocatalyst in the *Drosophila* system in comparison with TS-ΔIsPET. This observation indicates that LCC variants may represent promising candidates for further investigation in insect species relevant to waste bioconversion processes.

Our results contribute to the development of transgenic insect-based platforms as exploratory tools for plastic-associated waste processing. Recent considerations have highlighted that insect-mediated plastic degradation could result in secondary fragmentation and generation of micro- and nanoplastics, raising concerns about the environmental fate of plastic-derived particles [28]. In this scenario, intestinal expression of *in vitro*-evolved plastic-degrading enzymes in engineered insects may, in principle, complement these endogenous processes by promoting further depolymerisation of fragmented residues. Overall, this study represents an additional step toward possible strategies to mitigate the accumulation of persistent particulate plastic forms within controlled systems.

## 5. Materials and methods

### 5.1 Production of recombinant proteins in *E. coli*

The TS-ΔIsPET recombinant protein was expressed in the Origami 2 (DE3) *E. coli* strain and purified as reported in [44]. The synthetic genes (optimised for *E. coli* heterologous expression) encoding the wild-type and ΔLCC variants generated by site-directed mutagenesis lacking the native N-terminal signal peptide were cloned into the pET24b expression vector and expressed in the DE3 *E. coli* strain. The corresponding recombinant proteins were purified as reported in [45]. ΔLCC concentration was estimated based on the theoretical extinction coefficient at 280 nm of 38453 M^-1^cm^-1^.

### 5.2 Thermal stability of ΔLCC variants

The melting temperature (T_m_) for secondary structure of ΔLCC variants was determined by measuring the variation in ellipticity signal by circular dichroism (CD) at 222 nm during temperature ramps [75]. Proteins (0.1 mg/mL) were dissolved in 25 mM Tris-HCl, 200 mM NaCl, pH 8.0.

### 5.3 Enzymatic characterisation of recombinant proteins

The hydrolytic enzymatic activity on 1 mM p-nitrophenyl acetate (pNPA) was measured by recording the absorbance increase due to the product p-nitrophenolate at 415 nm (ε_415_ = 13.0 mM-^1^cm^-1^). The reaction was performed at 30 °C in 50 mM Na_2_HPO_4_, 100 mM NaCl, pH 7.0 with 1 mM pNPA at 30°C.

The activity of TS-ΔIsPET and ΔLCC (wild-type and variants) and the temperature dependence of the activity of the TA-ΔLCC variant on PET nanoparticles was assessed on PET nanoparticles by turbidimetric assay using 10-20 μg/mL enzyme and ∼0.1 mg/mL PET nanoparticle. The optical density variation at 600 nm was monitored for 10 min at the desired temperature after a 1 min preincubation before enzyme addition. The turbidity (OD_600_) was measured using a Jasco V-560 spectrophotometer (Jasco Inc.) and activity was determined from the slope of the linear region of the recorded curves [44].

PET nanoparticles (average diameter of 80 nm) were prepared from PET microplastics (diameter 300 μm; Goodfellow GmbH) using a precipitation and solvent evaporation technique [49].

### 5.4 Production of constructs for *D. melanogaster* S2R+ cell transfection and transgenic lines generation

The *TS-ΔIsPET* and *TA-ΔLCC* sequences were optimised for an efficient translation in *D. melanogaster* using the GenSmart™ Codon Optimization tool (https://www.genscript.com) and employed for *in silico* generating *Dm*-*TS-ΔIsPET* and *Dm*-*TA-ΔLCC* chimeric genes. Both genes included a translational enhancer sequence followed by a 51 bp coding sequence specifying the 17 aa signal peptide of a *D. melanogaster* chymotrypsin-like protein highly expressed in the insect midgut (Flybase ID: FBpp0077758) at their 5’ end, and a 6-His tag coding fragment at their 3’ end (Fig. 2A and Supplementary Fig. 3).

*Dm-TS-ΔIsPET* and *Dm-TA-ΔLCC* were synthetically produced and cloned in two different configurations: (i) as *5’-Kpn*I- *Hind*III-3’ segments into the pAc5-STABLE2-neo vector, deprived of both *Cherry* and *EGFP* marker genes, generating the *Dm-TS-ΔIsPET_pAc5*-STABLE2-neo and *Dm-TA-ΔLCC_pAc5*-STABLE2-neo constructs for cell transfection experiments [76] (Supplementary Fig. 4A, B) and (ii) as *5’-Kpn*I- *Xba*I-3’ fragments into the pJFRC81 vector, producing the *Dm-TS-ΔIsPET_pJFRC81* and *Dm-TA-ΔLCC_ pJFRC81* constructs for *D. melanogaster PhiC31* integrase-mediated transgenesis [77,78] (Supplementary Fig. 4C, D). The constructs were synthesised by GenScript Gene (https://www.genscript.com/).

### 5.5 Cell cultures and transfection

The *Drosophila* S2R+ cell line derived from a primary culture of 20-24 h old *D. melanogaster* embryos and was originally obtained from the *Drosophila* Genomics Resource Center (DGRC ID 150). S2R+ cells were cultured at 25 °C without CO_2_ in Schneider’s medium (Biowest) supplemented with 10 % heat-inactivated fetal bovine serum (FBS; Euroclone). S2R+ cells were transfected with the *Dm-TS-ΔIsPET_pAc5*-STABLE2-neo or *Dm-TA-ΔLCC_pAc5*-STABLE2-neo constructs, or with the pAc5-STABLE2-neo vector, which served as negative control. S2R+ cells were seeded in 6-well plates (8 × 10^5^ cell/well). After 24 h, they were transfected using 0.4 μg of each construct/well and the Effectene Transfection Reagent (Qiagen), according to the manufacturer’s instructions. Forty-eight h after transfection, stably transfected cells were selected using geneticin G418 (1 mg/mL; Invivogen). Non-transfected cells were grown in parallel as a transfection negative control. At different time points (2, 6, and 8 days post-transfection), cells were pelleted and stored at -80 °C in TRIzol™ Reagent (Thermo Fisher Scientific) for qPCR analyses. Supernatants from transfected and non-transfected samples were collected, supplemented with 5 % [v/v] glycerol (final concentration) and stored at -80 °C for Western blot analyses.

### 5.6 Cell viability assays

Cell viability was assessed at 2, 6, 8 and 10 days post-transfection using the Cell Counting Kit-8 (CCK-8) (Bimake). The CCK-8 reagent was added to each well according to the manufacturer’s instructions, and absorbance at 450 nm was measured using a microplate reader. Viability values were calculated relative to untreated control cells after background subtraction. For each time point, measurements were performed in triplicates.

### 5.7 Fly strains and maintenance

The *w^1118^* strain (# 5905) and the *daughterless-Gal4* (*daG4;* # 8641) driver were originally obtained from the Bloomington Drosophila Stock Center (Indiana University Bloomington). *Dm-TS-ΔIsPET-*O and *Dm-TA-ΔLCC-*O transgenic flies, carrying the *Dm-TS-ΔIsPET_pJFRC81* and *Dm-TA-ΔLCC_ pJFRC81* constructs, respectively, were obtained using the *PhiC31* integrase-mediated transgenesis systems by the Bestgene Inc (http://thebestgene.com; Chino Hills, CA 91709). The *Dm-TS-ΔIsPET-*O and *Dm-TA-ΔLCC-*O strains were maintained in heterozygosity with *CyO* and *TM3* balancer chromosomes, respectively. In adult flies, *CyO* and *TM3* balancers were identified by the presence of the dominant markers *Curly* (*Cy*; curly wings) and *Stubble* (*Sb;* short and thick bristles), respectively. Fly stocks were routinely maintained under 12 h light:12 h darkness (12:12 LD) regime at 23 °C, on a cornmeal standard diet (7.2 % [w/v] cornmeal, 7.9 % [w/v] sucrose, 5 % [w/v] inactive dried yeast, 0.85 % [w/v] agar, 0.3 % [v/v] propionic acid, and 0.27 % [w/v] nipagin).

### 5.8 Post-embryonic development, adult fertility, and survival

For the evaluation of *Dm-TS-ΔIsPET* and *Dm-TA-ΔLCC* overexpression effects on post-embryonic development, *daG4*>*Dm-TS-ΔIsPET-O* and *daG4*>*Dm-TA-ΔLCC-O* individuals were respectively compared to *daG4/CyO* and *daG4/TM3* controls, obtained by mating heterozygous *Dm-TS-ΔIsPET-O*/*CyO* or *Dm-TA-ΔLCC-O*/*TM3* flies to homozygous *daG4* individuals. For each cross, fertilised eggs were reared at 23 and 28 °C and monitored throughout development until eclosion. Once adult emergence occurred, flies were classified based on their genotype and sex.

To determine the fertility of *daG4*>*Dm-TS-ΔIsPET-O* and *daG4*>*Dm-TA-ΔLCC-O* flies, for each genotype ∼3-4-day old virgin females and males were mated in standard vials for 24 h. They were then subdivided into groups of 10 females and 10 males, and females were allowed to lay eggs on polystyrene plates (35 × 10 mm; Falcon), supplemented with fresh food, at 23 and 28 °C. After 24 h, the laid eggs were counted, and larval development was monitored for additional 48-72 h. Hatched larvae were counted and for each plate the percentage of developing larvae was calculated as the ratio between the number of hatched larvae and that of laid eggs.

Adult survival rate was assessed by comparing *daG4*>*Dm-TS-ΔIsPET-O* and *daG4*>*Dm-TA-ΔLCC-O* flies to their respective controls *daG4/+, Dm-TS-ΔIsPET-O/+,* and *Dm-TA-ΔLCC-O/+,* obtained by crossing individuals bearing either the *daG4* driver alone or the *Dm-TS-ΔIsPET* and *Dm-TA-ΔLCC* inserts alone to *w^1118^* flies. Adults of the same sex were maintained at a density of 10 individuals per vial at 23 and 28 °C. Flies were counted and transferred to fresh medium three times per week, with no anaesthesia.

### 5.9 *Dm-TS-ΔIsPET* and *Dm-TA-ΔLCC* mRNA expression analyses

*Dm-TS-ΔIsPET* and *Dm-TA-ΔLCC* mRNA expression analyses were performed via qPCR on transfected cells (for each construct, 3-5 replicates per time point), and on whole 3^rd^ instar larvae and dissected guts from *daG4*>*Dm-TS-ΔIsPET-O* and *daG4*>*Dm-TA-ΔLCC-O* genotypes as well as negative controls (*daG4/+, Dm-TS-ΔIsPET-O/+,* and *Dm-TA-ΔLCC-O/+*). Whole larvae were snap-frozen in liquid nitrogen, while guts were dissected in Phosphate-Buffered Saline (PBS, 137 mM NaCl, 2.7 mM KCl, 10 mM Na_2_HPO_4_/KH_2_PO_4_, pH 7.4) and placed in RNAlater^TM^ (Thermo Fisher Scientific). For each genotype, 3-5 replicates, each containing 10 larvae or 20 guts, were prepared and stored at -80 °C until use. Total RNA was extracted from cell and gut samples using Direct-zol RNA Miniprep kit (Zymo Research), and from whole larvae using TRIzol reagent (Thermo Fisher Scientific), following manufacturer’s instructions. Complementary DNAs (cDNAs) were synthesised using random primers (Promega) and the ImProm-II™ Reverse Transcriptase kit (Promega), accordingly to manufacturer’s instructions. For each sample, amplification was done in triplicate, using the GoTaq qPCR Master Mix (Promega) and a CFX96 Real-Time PCR Detection System (Bio-Rad), following the manufacturer’s instructions. Each qPCR reaction was performed in a 10 μL final volume, with 10 ng of cDNA and 0.2 μM of each primer. The amplification conditions were 95 °C 2 min, (15 sec at 95 °C, 1 min at 59 °C) for 40 cycles. Primers are reported in Supplementary Table 1. The standard housekeeping gene *rp49* (FlyBase ID: FBgn0002626) was used as an endogenous control. mRNA fold changes were calculated using the 2^-ΔΔCt^ method. For transfected S2R+ cells, the calibrators were the mean ΔCts of *Dm-TS-ΔIsPET* and *Dm-TA-ΔLCC* in the negative controls, transfected with the pAc5-STABLE2-neo vector. For transgenic larvae and guts, the calibrators were the mean ΔCts of *Dm-TS-ΔIsPET* and *Dm-TA-ΔLCC* in the *Dm-TS-ΔIsPET-O/+* and *Dm-TA-ΔLCC-O/+* controls, respectively.

### 5.10 Preparation of larval crude extracts

Samples from *daG4*>*Dm-TS-ΔIsPET-O* and *daG4*>*Dm-TA-ΔLCC-O* 3^rd^ instar larvae and negative controls (*daG4/+, Dm-TS-ΔIsPET-O/+* and *Dm-TA-ΔLCC-O/+*) (whole larva, gut, and carcass) were supplemented with 5 mg of the tyrosinase inhibitor phenylthiourea and homogenised with a mechanical homogeniser in RIPA buffer (50 mM Tris-HCl, 150 mM NaCl, 2 % [v/v] NP-40, 0.1 % [w/v] SDS, 5 mM EDTA and 1 μg/mL pepstatin, pH 8), using a 1:7 (mg/μL) ratio of larval tissue to RIPA buffer. Lysates were placed in 1.5 mL tubes (Eppendorf) and horizontally shaken on ice for 1 h at moderate speed. After centrifugation at 14,000 × g 15 min at 4 °C, supernatants were collected and stored at -80 °C for Western blot and enzyme activity analyses. Gut samples were mechanically lysed using the same procedure but using a motor-driven microtube pestle in 1.5 mL tubes (Eppendorf) on ice.

### 5.11 Enzymatic N-deglycosylation of Dm-TS-ΔIsPET, and Dm-TA-ΔLCC recombinant proteins

Crude extracts of larvae were subjected to enzymatic digestion to remove oligosaccharides using peptide N-glycosidase F (PNGase F; New England Biolabs) according to the supplier’s instructions. Briefly, 9 μL of sample, containing ∼0.5 mg of proteins, were denatured for 10 min at 99 °C in glycoprotein-denaturing buffer at a final volume of 10 μL. After cooling, samples were treated with 500 U of PNGase F for 4 h at 37 °C. The reaction was stopped by adding SDS-PAGE sample buffer and incubating at 99 °C for 10 min. Samples were stored at -20 °C until Western blot analysis.

### 5.12 Production and validation of custom antibodies and Western blot of Dm*-*TS-ΔIsPET and Dm*-*TA-ΔLCC

Rabbit polyclonal antibodies raised against TS-ΔIsPET and TA-ΔLCC (Davids Biotechnologie, Regensburg, Germany) were purified from rabbit antiserum by ammonium sulphate precipitation at 50 % saturation. Precipitated antibodies were collected by centrifugation at ∼41,000 g for 1 h at 4 °C, resuspended in 20 mM potassium phosphate buffer (pH 8.0) and dialysed against the same buffer. The specificity and sensitivity of the custom antibodies were validated by Western blot analysis (Supplementary Fig. 12).

Western blot analyses were performed on supernatants of cell cultures (after centrifugation at 14,000 × g for 15 min at 4 °C) and of larval crude extracts. Protein content was quantified using the Bradford Protein assay. Samples were supplemented with 4× SDS loading buffer (Thermo Fisher Scientific) and 0.1 M dithiothreitol (Merck), denatured at 95 °C for 5 min and resolved onto 12 % [w/v] acrylamide gels, either custom-made or precast (NuPAGE™ Bis-Tris Mini Protein Gels; Thermo Fisher Scientific). After separation, proteins were transferred onto a nitrocellulose membrane (GE Healthcare). Membranes were blocked for 1 h at room temperature (RT) with 5 % [w/v] skimmed dry milk in Tris-Buffered Saline (20 mM Tris, pH 7.4; 150 mM NaCl) containing 0.05 % [v/v] Tween (TBST), and incubated overnight at 4 °C with the following primary antibodies: i) mouse anti-His antibody (1:5000; Thermo Fisher Scientific) and ii) custom rabbit polyclonal antibodies raised against TS-ΔIsPET and TA-ΔLCC (both used at 0.2 mg/mL final concentration) (Davids Biotechnologie, Regensburg, Germany). After washing in TBST, membranes were incubated with the appropriate secondary Horseradish Peroxidase (HRP)-anti IgG antibodies for 2 h at RT. Chemiluminescence signals were visualised using custom-made or commercial enhanced chemiluminescence (ECL) buffers (ECL Plus WB Detection Kit, GE Healthcare). Prestained Protein SHARPMASS™ V plus (Euroclone) or PageRulerTM (Prestained Protein Ladder, Thermo Fisher Scientific) were used as protein molecular weight markers.

### 5.13 Enzymatic degradation of PET

The biodegradation reactions of PET nanoparticles were set up adding 0.4 mL of 0.6 mg/mL PET nanoparticle solution to 1 mL final volume of reaction solution (50 mM Na_2_HPO_4_, pH 9, 100 mM NaCl). Reaction was started by the addition of 30 μL of larval extracts or 0.6 μg recombinant microbial TA-ΔLCC (positive control). The reaction was incubated at 55 °C under rotatory agitation. The combined concentration of the soluble aromatic products MHET, TPA, and bis(2-hydroxyethyl) terephthalate (BHET) was determined by recording the absorbance spectrum of the reaction mixture (ε_240_ = 13.8 mM^-1^cm^-1^) and by HPLC separation [49]. For HPLC analysis, 33 μL of the reaction mixture was diluted with 67 μL of the mobile phase (60 % of 0.1 % formic acid, and 40 % of methanol). 20 μL of each sample were injected into the HPLC column Symmetry^®^ C18, 5 μm (Waters™) and products were detected using an UV/Vis detector at 240 nm. Separation was performed at 35 °C. The concentration of TPA (retention time 9.5 min) and MHET (retention time 10.5 min) was determined from the areas of the corresponding peaks using the following equations: [TPA] = Peak area/2.32 × 10⁷ (range: 0.001 - 0.1 mM) [MHET] = Peak area + 9799/3.00 × 10^7^ (range: 0.025 - 0.075 mM)

### 5.14 Transmission Electron Microscopy (TEM), whole-mount immunostaining, immunogold labelling, and APN and ALP enzymatic activities of larval guts

Gut samples were obtained from *daG4*>*Dm-TA-ΔLCC-O* 3^rd^ instar larvae and negative controls (*daG4/+,* and *Dm-TA-ΔLCC-O/+*).

#### 5.14.1 TEM analysis

Midgut samples were dissected in PBS and fixed overnight in 2 % [v/v] glutaraldehyde in 0.1 M Na-cacodylate buffer (pH 7.4) at 4 °C. Specimens were post-fixed with 1 % osmium tetroxide (in 0.1 M Na-cacodylate buffer, pH 7.4) for 1 h at RT and subsequently dehydrated in ascending ethanol series. Finally, they were embedded in Epon/Araldite 812 mixture resin. 60-nm-thick sections were obtained with a Leica Reichert Ultracut S (Leica) and stained with lead citrate and uranyl acetate before analysis with a JEM-1400Flash transmission electron microscope (Jeol) equipped with a Morada digital camera (Olympus).

#### 5.14.2 Whole-mount immunostaining

Guts were dissected in PBS, fixed in 4 % [w/v] paraformaldehyde (in PBS) for 20 min at RT, and permeabilised in PBS with 0.5 % [v/v] Triton X-100 for 10 min. Samples were then pre-incubated for 1 h with 5 % goat serum (Thermo Fisher Scientific) in 0.1 % [v/v] Triton X-100, before incubation with anti-TA-ΔLCC antibody (0.2 mg/mL in 5 % goat serum, 0.5 % [v/v] Triton X-100) overnight at 4 °C. After 3 washes of 10 min with 0.5 % [v/v] Triton X-100, specimens were incubated for 2 h at RT with an anti-rabbit FITC-conjugated antibody (1:200 in 0.5 % [v/v] Triton X-100; Jackson ImmunoResearch). After 3 washes with 0.5 % [v/v] Triton X-100 (10 min each) and 2 with PBS (5 min each), samples were incubated with 4′,6-diamidino-2-phenylindole (DAPI) (100 ng/mL in PBS) for nuclear staining. Specimens were mounted on glass slides with Citifluor (Citifluor Ltd) and analysed with an Eclipse Ni-U microscope equipped with a DS-SM-L1 digital camera (Nikon), and with a Leica Stellaris 5 confocal microscope (Leica). Negative controls were performed by omitting the primary antibody to assess non-specific binding (Supplementary Fig. 13).

#### 5.14.3 Immunogold labelling

Gut cross-sections were obtained as reported above for TEM analyses and treated with 3 % NaOH in ethanol (w/v) to remove the resin. Sections were then incubated with 2 % [w/v] Bovine Serum Albumin (BSA), 0.1 % [v/v] Tween 20 in PBS for 30 min and then with anti-TA-ΔLCC antibody (1.3 mg/mL) for 1 h. After 2 washes with 2 % BSA, 0.1 % v/v Tween 20 in PBS for 10 min each, specimens were incubated with 10 nm gold-conjugated goat anti-rabbit antibody (GE Healthcare Life Sciences; dilution 1:50) for 1 h. Afterwards, sections were post-fixed with 0.5 % v/v glutaraldehyde in PBS for 5 min, counterstained with uranyl acetate, and observed under a JEM-1400Flash transmission electron microscope (Jeol). Negative controls were performed by omitting the primary antibody to assess non-specific binding (Supplementary Fig. 14).

#### 5.14.4 APN and ALP enzymatic activities

Midguts were isolated in PBS, blotted on filter paper, and frozen in dry ice. Pools of 20-30 midguts were stored at -80 °C until use. After thawing on ice, midgut samples were homogenised with a microtube pestle in 100 mM mannitol, 10 mM Hepes-Tris, pH 7.2 (1 mL/100 mg tissue). Protein concentration was determined by Bradford Protein assay using BSA as standard. The activity of aminopeptidase N (EC 3.4.11.2) (APN) and alkaline phosphatase (EC 3.1.3.1) (ALP) in the homogenate was determined as previously described [61], measuring spectrophotometrically the release of p-nitroanilide (pNA) from L-leucine p-nitroanilide and p-nitrophenol (pNP) from p-nitrophenyl phosphate, respectively. One unit (U) of enzyme activity was defined as the amount of enzyme that releases 1 mmol of pNA or pNP per min per mg of proteins. Assays were performed under conditions in which product formation was linearly associated with enzyme concentration.

### 5.15 PET nanoparticle feeding assays

#### 5.15.1 Umbelliferone-labelled PET nanoparticle feeding and confocal imaging

PET nanoparticles were labelled with the blue fluorophore umbelliferone (UMB-PET), which was incorporated during nanoparticle preparation at 0.1 % relative to PET. UMB-PET nanoparticles were added to an instant medium (24 % [w/v] mashed potato flakes, 8 % [w/v] sucrose, 2.4 % [w/v] inactive dry yeast, 0.3 % [v/v] propionic acid, and 0.27 % [w/v] nipagin in distilled water) at a final concentration of 0.05 % (500 µg/g medium). Instant medium without UMB-PET nanoparticles represented the control food.

*w¹¹¹⁸* embryos were reared on UMB-PET-supplemented or PET-free food at 28 °C until the 3^rd^ larval stage. Larval guts were dissected in PBS, fixed in 4 % [w/v] paraformaldehyde for 20 min at RT, washed in PBS, and mounted in VECTASHIELD mounting medium (VECTOR Laboratories) supplemented with propidium iodide (PI; 1 µg/mL; Sigma). Samples were imaged using a Leica Stellaris 8 confocal microscope (Leica) under identical acquisition settings.

#### 5.15.2 PET nanoparticle exposure assays

*daG4*>*Dm-TA-ΔLCC-O* 24 h-old embryos and the corresponding negative controls (*Dm-TA-ΔLCC-O/*+ and *daG4/+*) were transferred to instant medium supplemented with 0.05 % PET nanoparticles (500 µg/g medium). Control groups were reared on the same instant medium without PET nanoparticles. For each genotype and condition, four replicates with 45–80 embryos each, were monitored at 28 °C throughout post-embryonic development until adult eclosion, as described above. To assess fertility, *daG4*>*Dm-TA-ΔLCC-O* adults reared in the presence or absence of PET nanoparticles were crossed on standard medium at 28 °C. For each condition, three replicates (67–170 embryos each) were monitored until the 3^rd^ larval instar, as described above.

### 5.16 Statistical analyses

Fisher’s exact test was used to evaluate post-embryonic development data. Adult survival was analysed using the Mantel-Cox test. The normality of qPCR data, APN and ALP activity measurements, egg-to-adult viability and adult fertility data from PET nanoparticle feeding assays was assessed by Lilliefors (Kolmogorov-Smirnov) and/or Shapiro-Wilk tests. Data not following a normal distribution were analysed with the non-parametric Kruskal-Wallis test, followed by Dunn’s *post hoc* test. Normally distributed data were subjected to either ordinary one-way ANOVA, followed by Holm-Šídák’s multiple comparisons test, or the Brown-Forsythe test (for samples with significantly different standard deviations), followed by Dunnett’s *post hoc* test. Unpaired two-tailed t-test, two-way ANOVA, or two-way repeated-measures (RM) ANOVA were used when appropriate. Analyses and graphs were performed using GraphPad Prism 10.4.1 and KaleidaGraph 4.0.

## Author contributions

F.S., G.M., and G.T. conceived the study. F.S., G.M., V.P., A.G., and G.T. designed the experiments. V.P., F.B., D.B., Sa.C., C.F., M.B., and D.R. performed the experiments. V.P., F.B., D.B., M.B., E.C.-C., D.R., Si.C., F.S. and G.M. analysed the data. F.S., G.M., A.G., Si.C., M.C., and G.T. supervised the project. G.M., F.S., D.B., Si.C., and G.T. wrote the manuscript. F.S., G.M., A.G., Si.C., M.C., and G.T. edited the manuscript. All authors read and approved the final manuscript.

## Data availability

The data supporting the findings of this study are available from the corresponding authors upon request.

## Acknowledgements

We thank the DiBio fly facility (https://dibio-fly-facility.netlify.app/) at the Department of Biology, Università degli Studi di Padova for *Drosophila* lines maintenance. We are grateful to Davide Tessaro (Politecnico di Milano) for the production of PET nanoparticles and Nicolò Picchio for support in HLPC analysis. Scientific support from CRIETT center of University of Insubria (instrument codes MIC01, MIC03, and MIC06) is greatly acknowledged. We thank Loredano Pollegioni for insightful comments on the biochemistry sections.

The project was supported through funding by PRIN (Research Projects of National Interest) Program 2020 - Italian Ministry of Education, Universities and Research (MUR) (Prot. 2020ENH3NZ) to F.S., G.T., and Si.C.; Consorzio Italiano per le Biotecnologie CIB (G.M. and G.T.); and Fondo di Ateneo per la Ricerca (G.M. and G.T.).

