## Supplementary material for "*In vivo* expression of engineered IsPETase and LCC variants in *Drosophila melanogaster* reveals host tolerance and differential catalytic activity"

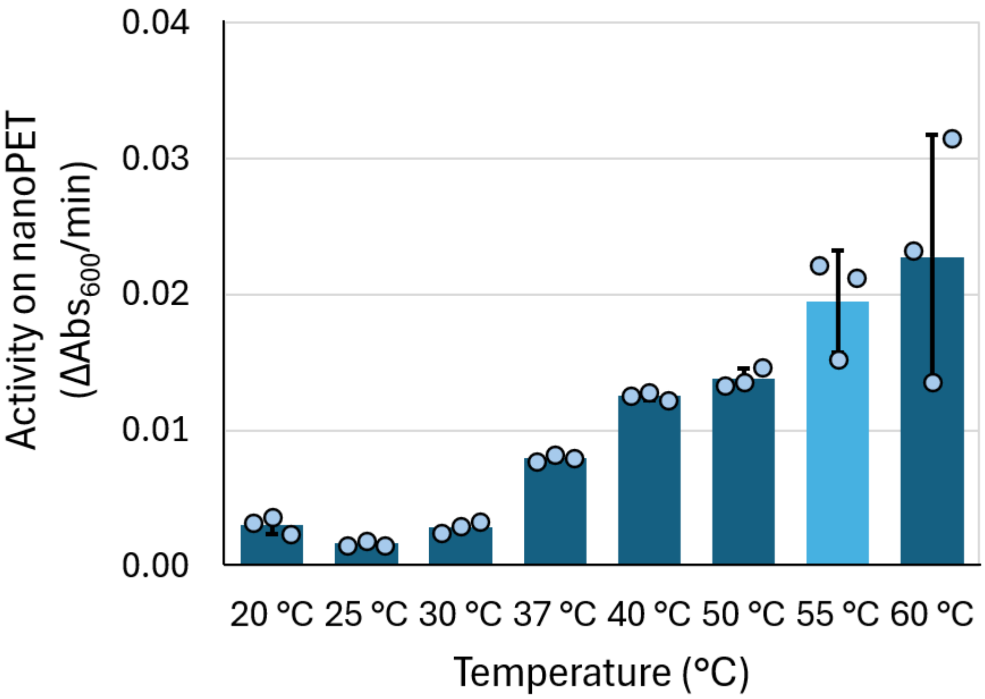


**Supplementary Figure 1.** Effect of reaction temperature on the activity of TA-ΔLCC. Reactions were performed in 50 mM $Na_2HPO_4$, 100 mM NaCl, pH 9.0, on 0.1 mg/mL PET nanoparticles. TA-ΔLCC concentration: 10 µg/mL. ΔAbs/min values are shown as absolute values. Assays were performed in triplicate (n = 3) at each temperature.

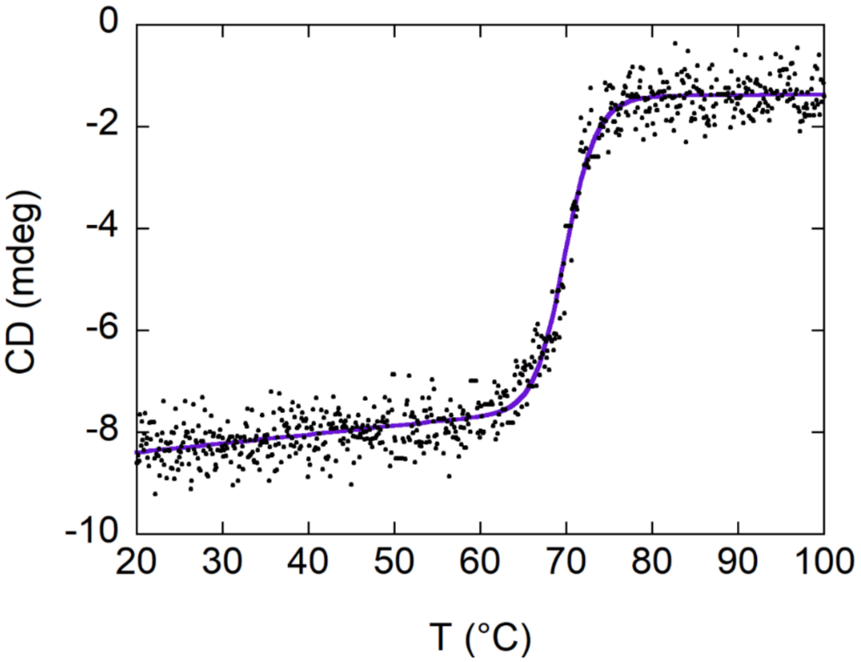


**Supplementary Figure 2.** Circular dichroism (CD) thermal denaturation profile of TA-ΔLCC variant. Denaturation was monitored by CD spectrometry at 222 nm. The curve was fitted to a sigmoidal function providing a $T_m$ of 69.9 ± 0.1 °C.

**A *Dm-TS-ΔIsPET***

```
1   atg aag cag ttc gcc gtc atc ttt gct ttg gcg ctg gcc agc gtg tcc gca caa acc aat 60
     M   K   Q   F   A   V   I   F   A   L   A   L   A   S   V   S   A   Q   T   N
61  cct tat gcg aga ggt ccc aat ccg aca gct gcc agt ttg gaa gca tca gcg ggt cca ttc 120
     P   Y   A   R   G   P   N   P   T   A   A   S   L   E   A   S   A   G   P   F
121 acg gtc cgg agc ttc acc gta tcg cgt cca tcc gga tat ggc gcc ggc acc gtc tac tac 180
     T   V   R   S   F   T   V   S   R   P   S   G   Y   G   A   G   T   V   Y   Y
181 cct aca aat gct ggc gga act gtt gga gcc atc gcc att gta ccc ggc tac acg gct cgc 240
     P   T   N   A   G   G   T   V   G   A   I   A   I   V   P   G   Y   T   A   R
241 caa tcc tcc atc aag tgg tgg ggc cca cgt ctg gcc tcg cac gga ttt gtg gtg atc acc 300
     Q   S   S   I   K   W   W   G   P   R   L   A   S   H   G   F   V   V   I   T
301 att gat acg aac tcg acg ctc gat cag ccg gag agt cgg tcg tca cag cag atg gca gca 360
     I   D   T   N   S   T   L   D   Q   P   E   S   R   S   S   Q   Q   M   A   A
361 ctg cgc cag gtg gct tcg ctg aat ggc aca agt tcc tcc ccg ata tat ggc aag gtg gac 420
     L   R   Q   V   A   S   L   N   G   T   S   S   S   P   I   Y   G   K   V   D
421 acc gca cgc atg ggt gtg atg gga cac tcg atg ggc ggg ggt gga tca ctt ata agc gcg 480
     T   A   R   M   G   V   M   G   H   S   M   G   G   G   G   S   L   I   S   A
481 gcc aac aat ccc agc ttg aaa gca gct gct ccg cag gca ccc tgg cac agc agc acc aac 540
     A   N   N   P   S   L   K   A   A   A   P   Q   A   P   W   H   S   S   T   N
541 ttc agc tct gtg acc gtt cca aca cta atc ttc gcg tgc gaa aac gat tct att gcc ccg 600
     F   S   S   V   T   V   P   T   L   I   F   A   C   E   N   D   S   I   A   P
601 gtc aat agt agt gcc ctg ccc atc tac gat tcc atg tcc agg aat gcc aag cag ttc ctg 660
     V   N   S   S   A   L   P   I   Y   D   S   M   S   R   N   A   K   Q   F   L
661 gag atc aac ggt ggc agc cat gcc tgt gcc aat tcc ggc aac agc aac caa gcg ctc att 720
     E   I   N   G   G   S   H   A   C   A   N   S   G   N   S   N   Q   A   L   I
721 gga aaa aag ggc gtg gcc tgg atg aag cgc ttc atg gac aac gac act cga tac tcc acc 780
     G   K   K   G   V   A   W   M   K   R   F   M   D   N   D   T   R   Y   S   T
781 ttt gcc tgc gag aac ccc aat agc act cga gtt tcg gac ttt agg acg gcg aac tgc tct 840
     F   A   C   E   N   P   N   S   T   R   V   S   D   F   R   T   A   N   C   S
841 cac cac cat cat cac cat 858
     H   H   H   H   H   H
```

**B *Dm-TA-ΔLCC***

```
  1 atg aag cag ttc gcc gtc atc ttt gct ttg gcg ctg gcc agc gtg tcc gca caa agc aat 60
     M   K   Q   F   A   V   I   F   A   L   A   L   A   S   V   S   A   Q   S   N
 61 cca tat cag agg gga ccc aat ccc act aga tca gcg ctc acg gcg gat ggc cca ttt tcg 120
     P   Y   Q   R   G   P   N   P   T   R   S   A   L   T   A   D   G   P   F   S
121 gtg gcc acc tac act gtc tct agg ctt tca gta tcc ggc ttt gga ggt ggc gtt atc tac 180
     V   A   T   Y   T   V   S   R   L   S   V   S   G   F   G   G   G   V   I   Y
181 tac cca aca ggc acc agc cta acc ttt ggc ggc ata gcc atg tcc cca gga tac acg gca 240
     Y   P   T   G   T   S   L   T   F   G   G   I   A   M   S   P   G   Y   T   A
241 gac gct tct tcg ctg gcc tgg ctg ggt cgt cgg cta gcc tcc cat ggc ttc gtg gtg ttg 300
     D   A   S   S   L   A   W   L   G   R   R   L   A   S   H   G   F   V   V   L
301 gtg atc aac aca aat tcg cgc ttc gat tat ccg gat agt cgt gcc tcg cag ctg tcg gca 360
     V   I   N   T   N   S   R   F   D   Y   P   D   S   R   A   S   Q   L   S   A
361 gct ctt aac tac ctg cgc acg tcc agt cca tca gct gtt cga gca cga tta gat gcc aat 420
     A   L   N   Y   L   R   T   S   S   P   S   A   V   R   A   R   L   D   A   N
421 cgc ttg gcc gtc gcg ggt cat agc atg ggt gga gga ggg acg tta aga ata gcc gag caa 480
     R   L   A   V   A   G   H   S   M   G   G   G   G   T   L   R   I   A   E   Q
481 aat ccg agc ctg aag gcc gcg gtg ccg ttg acg ccc tgg cac acc gac aag acc ttc aac 540
     N   P   S   L   K   A   A   V   P   L   T   P   W   H   T   D   K   T   F   N
541 act tcc gta cct gtc ttg att gtg ggc gct gaa gcc gat aca gtt gca ccg gtt agt caa 600
     T   S   V   P   V   L   I   V   G   A   E   A   D   T   V   A   P   V   S   Q
601 cat gcc att ccc ttc tac cag aac ctg ccc agc acc act cct aaa gtc tat gtg gag ctg 660
     H   A   I   P   F   Y   Q   N   L   P   S   T   T   P   K   V   Y   V   E   L
661 gac aat gcc agc cac aca gct ccc aac agc aac aac gcg gcc atc agt gtc tac acc att 720
     D   N   A   S   H   T   A   P   N   S   N   N   A   A   I   S   V   Y   T   I
721 tcc tgg atg aag ctc tgg gtg gat aac gac aca cgg tat cgc cag ttc ctg tgc aat gtg 780
     S   W   M   K   L   W   V   D   N   D   T   R   Y   R   Q   F   L   C   N   V
781 aac gat ccg gct ctc tcc gac ttc cgc acc aac aat cga cac gcc cag cac cac cat cat 840
     N   D   P   A   L   S   D   F   R   T   N   N   R   H   A   Q   H   H   H   H
841 cac cac 846
     H   H
```

**Supplementary Figure 3. Nucleotide and amino acid sequences of *Dm-TS-ΔIsPET* (A, 858 nt) and *Dm-TA-ΔLCC* (B, 846 nt) recombinant genes.**

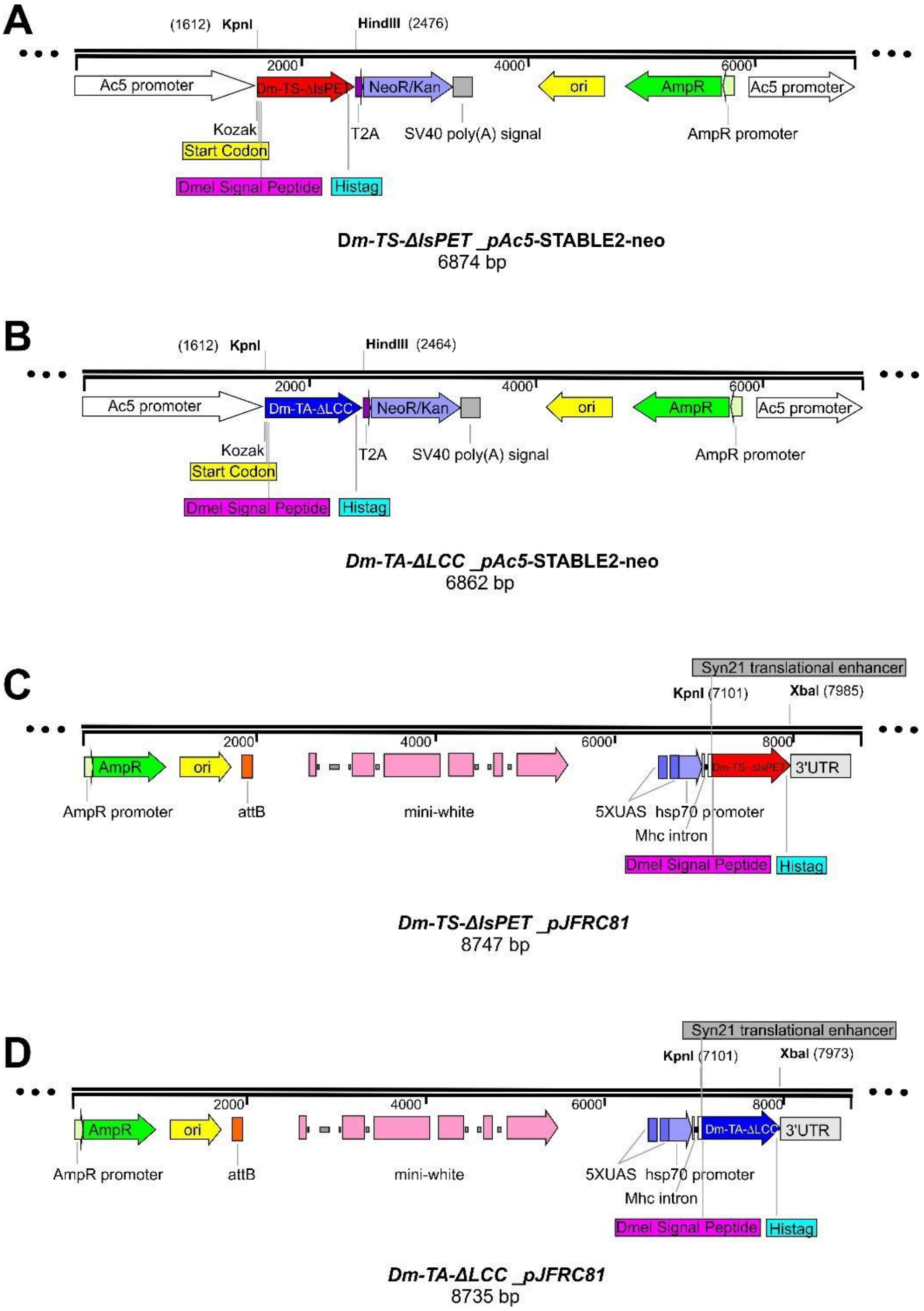


**Supplementary Figure 4. Schematic representations of the constructs used for cell transfection experiments and *D. melanogaster* transgenic generation. A, B** *Dm-TS-ΔIsPET* (red) and *Dm-TA-ΔLCC* (blue) DNAs cloned as *5'-Kpn*I-*Hind*III-3' segments into the pAc5-STABLE2-neo vector, deprived of *Cherry* and *EGFP* marker genes, in *Dm-TS-ΔIsPET_pAc5*-STABLE2-neo (**A**) and *Dm-TA-ΔLCC_pAc5*-STABLE2-neo (**B**) constructs for cell transfection experiments. **C, D** *Dm-TS-ΔIsPET* (red) and *Dm-TA-ΔLCC* (blue) DNAs cloned as *5'-Kpn*I- *Xba*I-3' fragments into the pJFRC81 vector, in *Dm-TS-ΔIsPET_pJFRC81* (**C**) and *Dm-TA-ΔLCC_pJFRC81* (**D**) constructs for *D. melanogaster PhiC31* integrase-mediated transgenesis.

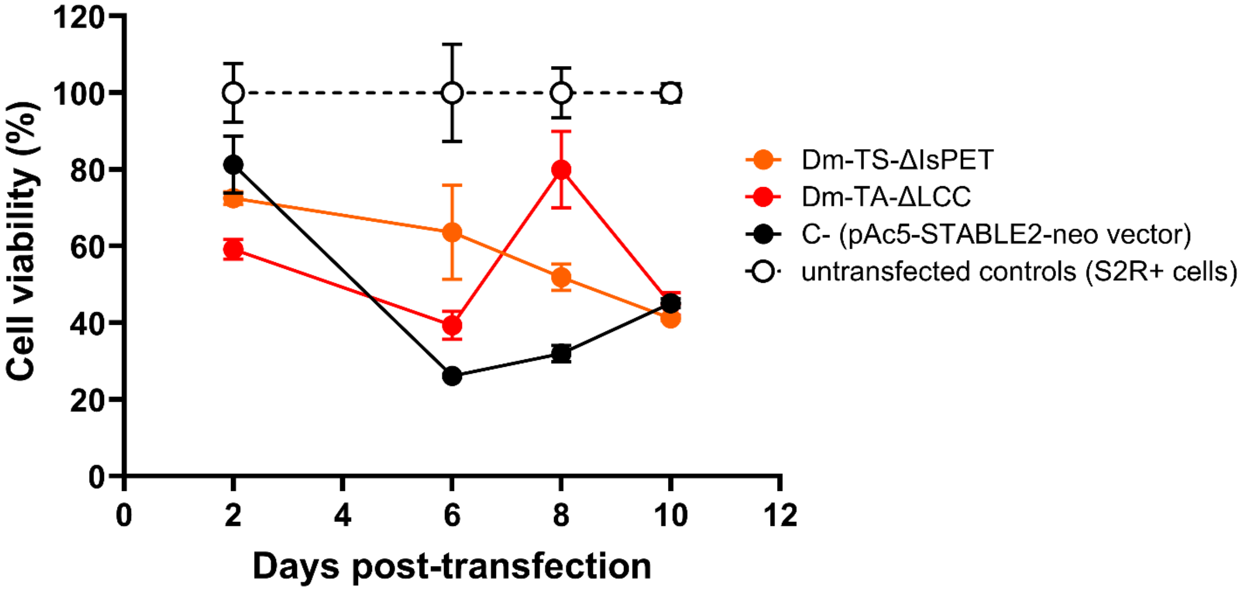


**Supplementary Figure 5. Cell viability assessed by CCK-8 assay.** Cell viability (mean ± SEM, 3 replicates per condition) of *Dm-TS-ΔIsPET-* (orange), *Dm-TA-ΔLCC-* (red) and pAc5-STABLE2-neo vector-transfected S2R+ cells (C-; black) evaluated at 2, 6, 8 and 10 days post-transfection. Viability of untransfected S2R+ cells (dashed line) is shown as a transfection-negative control. Two-way ANOVA for repeated measures (RM ANOVA) performed on transfected conditions revealed a significant effect of time ($F_{1.932, 15.45} = 5.319$; $P = 0.02$), while no significant differences were detected between transfected cell lines ($F_{2, 6} = 3.535$; $P = 0.096$).

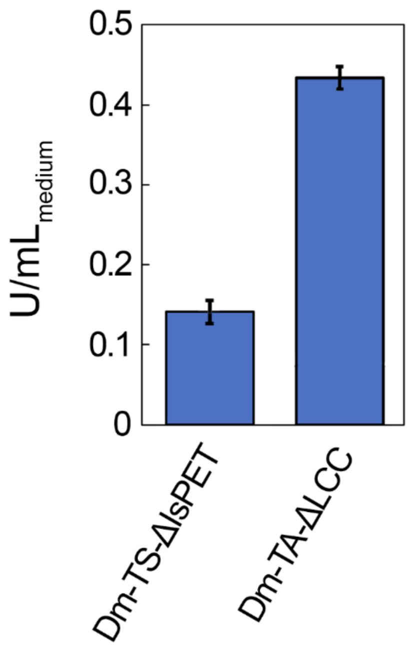


**Supplementary Figure 6. Activity of Dm-TS-ΔIsPET and Dm-TA-ΔLCC secreted from *Dm-TS-ΔIsPET*- and *Dm-TA-ΔLCC*-transfected S2R+ *D. melanogaster* cells.** Supernatants from transfected cell cultures were collected 6 days after transfection and assayed for esterase activity. For each assay, 100 µL of cell culture supernatant containing the enzyme of interest were analysed. The absorbance at 415 nm was recorded for 60 s. Activity data were subtracted by the corresponding blank (supernatants of cells transfected with the mock vector). Assays were performed in triplicate (n = 3).

**A**

| Parental lines | E | L3 | P | F1 progeny Adults | ♀ | ♂ | Tot | P (Fisher's exact test) |
|---|---|---|---|---|---|---|---|---|
| *daG4 X Dm-TS-ΔIsPET-O/CyO* | 148 | 134 | 132 | *daG4>Dm-TS-ΔIsPET-O* | 34 | 35 | 69 | P = 0.45, ns |
| | | | | *daG4>CyO* | 19 | 28 | 47 | |
| *daG4 X Dm-TA-ΔLCC-O/TM3* | 310 | 297 | 274 | *daG4>Dm-TA-ΔLCC-O* | 50 | 58 | 108 | P > 0.99, ns |
| | | | | *daG4>TM3* | 66 | 79 | 145 | |

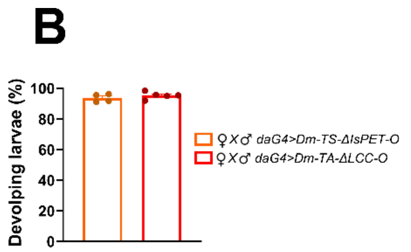


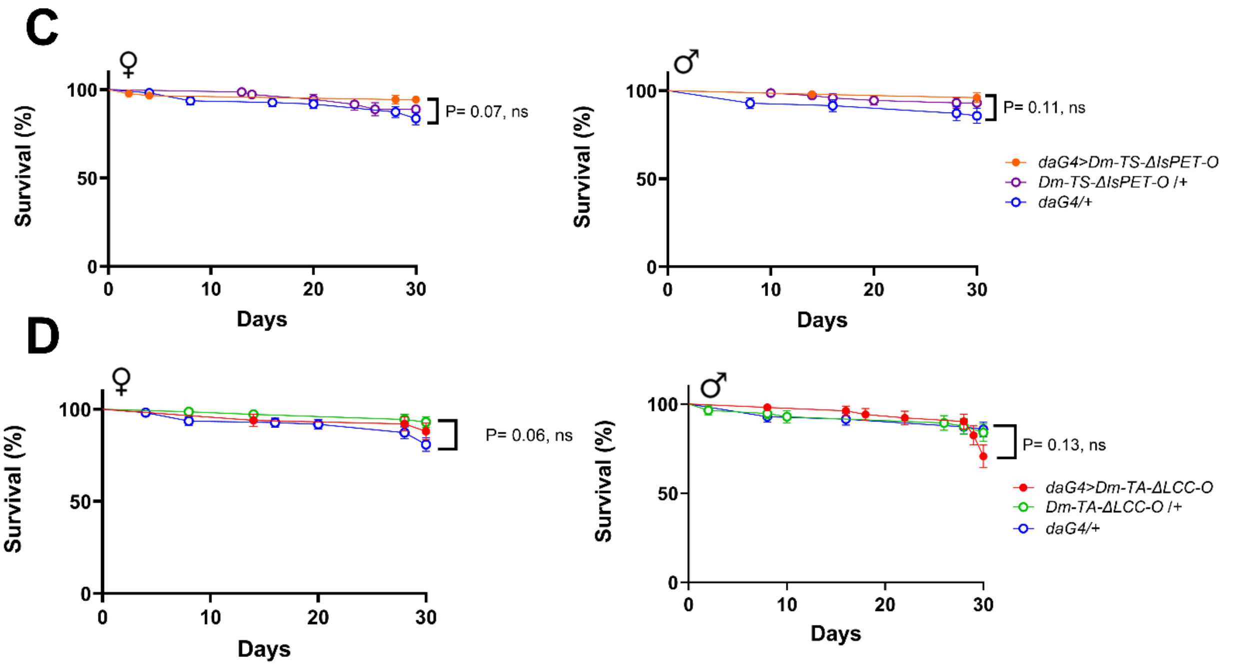


**Supplementary Figure 7. Effects of ubiquitous expression of *Dm-TS-ΔIsPET* and *Dm-TA-ΔLCC* on post-embryonic development, adult fertility, and survival at 28 °C. A** F1 progeny obtained by mating homozygous *daG4* with heterozygous *Dm-TS-ΔIsPET-O*/*CyO* flies and by mating homozygous *daG4* with heterozygous *Dm-TA-ΔLCC-O*/*TM3* flies. For each cross, total numbers of embryos (E), 3$^{rd}$ instar larvae (L3), and pupae (P) are reported. F1 adult females and males are reported based on their genotype. No significant differences were detected in Fisher's exact test between female and male *daG4>Dm-TS-ΔIsPET-O* flies and female and male *daG4/CyO* controls (P> 0.45, ns), as well as between female and male *daG4>Dm-TA-ΔLCC-O* flies and female and male *daG4/TM3* controls (P>0.99, ns). **B** Percentage (mean ± SEM) of developing larvae from embryos laid by *daG4>Dm-TS-ΔIsPET-O* (orange) and *daG4>Dm-TA-ΔLCC-O* (red) flies. Number of analysed embryos: *daG4>Dm-TS-ΔIsPET-O*: 643 [4 replicates (dots)]; *daG4>Dm-TA-ΔLCC-O*: 886 [5 replicates (dots)]. **C, D** Survival curves (mean % ± SEM) of *daG4>Dm-TS-ΔIsPET-O* (**C**) and *daG4> Dm-TA-ΔLCC-O* (**D**) flies, compared to their respective negative controls (*daG4/+*, and *Dm-TS-ΔIsPET-O/+* or *Dm-TA-ΔLCC-O/+*). No significant differences in Mantel-Cox test were detected among *daG4>Dm-TS-ΔIsPET-O, daG4/+*, and *Dm-TS-ΔIsPET-O/+* flies in both females (left) (P= 0.07, ns) and males (right) (P= 0.11, ns) in (**C**), and among *daG4>Dm-TA-ΔLCC-O, daG4/+*, and *Dm-TA-ΔLCC-O/+* flies in both females (left) (P= 0.06, ns) and males (right) (P= 0.13, ns) in (**D**). Number of analysed females and males, respectively: *daG4>Dm-TS-ΔIsPET-O:* 89, 50; *daG4/+*: 110, 70; *Dm-TS-ΔIsPET-O/+*: 72*,* 72; *daG4> Dm-TA-ΔLCC-O*: 50*,* 51; *Dm-TA-ΔLCC-O /+*: 72, 56.

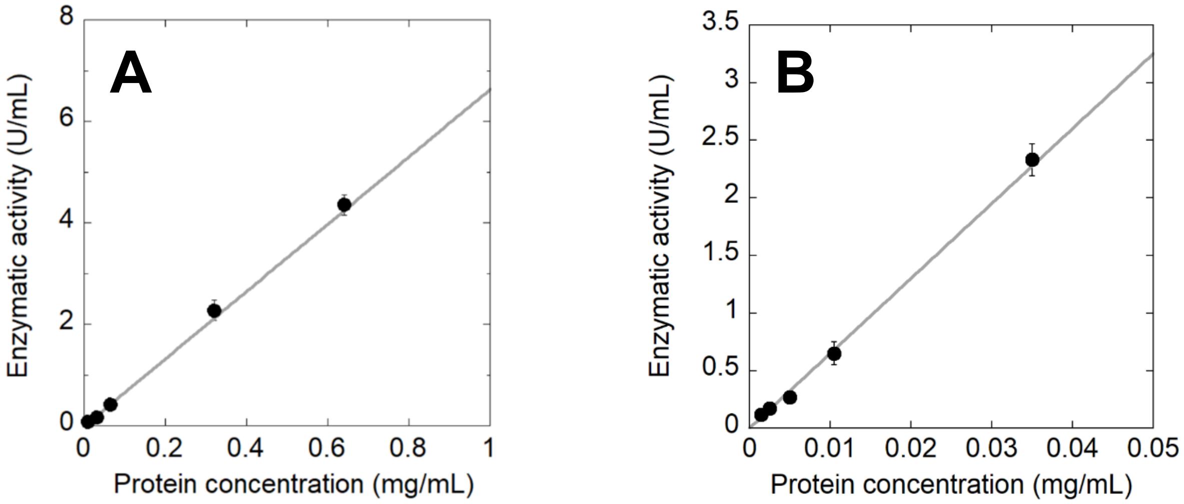


**Supplementary Figure 8. Calibration curves for the determination of amount of expressed PET hydrolytic enzymes**. **A** Enzymatic units of TS-ΔIsPET *vs.* enzyme concentration (µg/mL). **B** Enzymatic units of TA-ΔLCC *vs.* enzyme concentration (µg/mL). Assay was performed in 50 mM $Na_2HPO_4$, 100 mM NaCl, pH 7.0 with 1 mM pNPA at 30 °C (n=3).

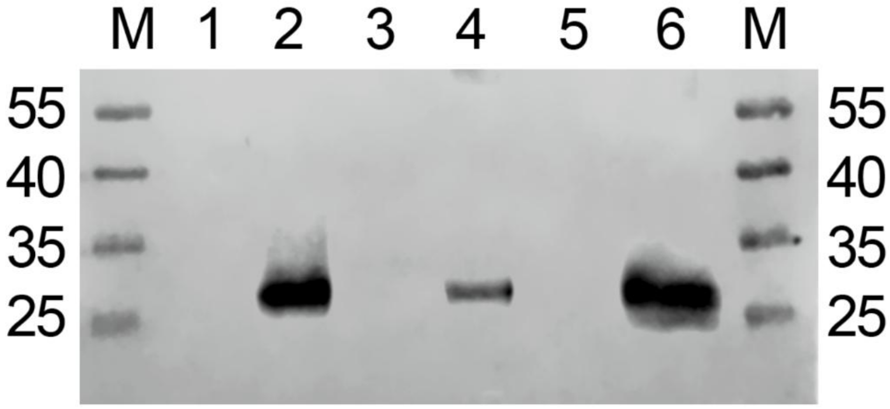


**Supplementary Figure 9. Identification of recombinant Dm-TA-ΔLCC in transgenic larvae.** Western blots of larval extracts using anti-TA-ΔLCC antibodies. M: molecular mass standards; lane 2: crude extract from 1 mg of whole larvae from *daG4>Dm-TA-ΔLCC-O* line (~0.2 mg proteins); lane 4: crude extract from 3 mg of carcasses from *daG4>Dm-TA-ΔLCC-O* line (~1.0 mg proteins); lane 6: crude extract from 3 mg of guts from *daG4>Dm-TA-ΔLCC-O* line (~0.3 mg proteins); lanes 1, 3, and 5: corresponding crude extracts (same amount of biomass of lanes 2, 4 and 6) from the corresponding negative controls from *Dm-TA-ΔLCC-O*/+ line.

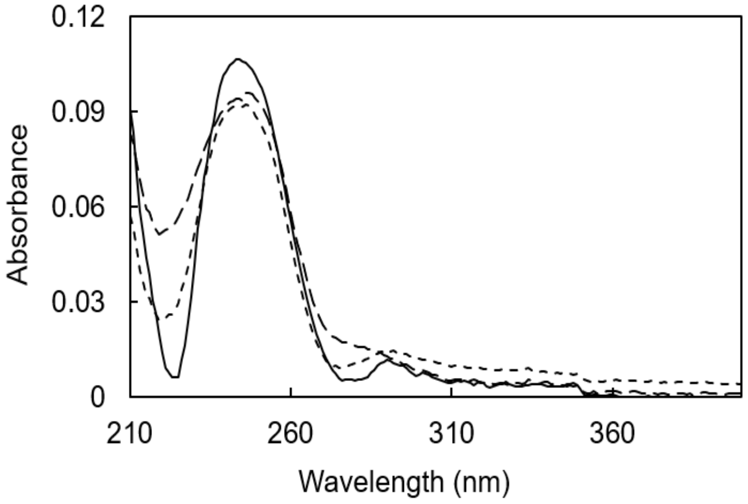


**Supplementary Figure 10. Depolymerisation of PET nanoparticles by extracts from Dm-TA-ΔLCC expressing larvae (*daG4>Dm-TA-ΔLCC-O*).** Differential absorbance spectrum of 3 independent reaction mixtures after 48 h of incubation (i.e., the spectrum shown in figure has been obtained by subtraction of the absorbance spectrum of the mixture at the beginning of the reaction). Spectra data points have been smoothed (window size 5 nm).

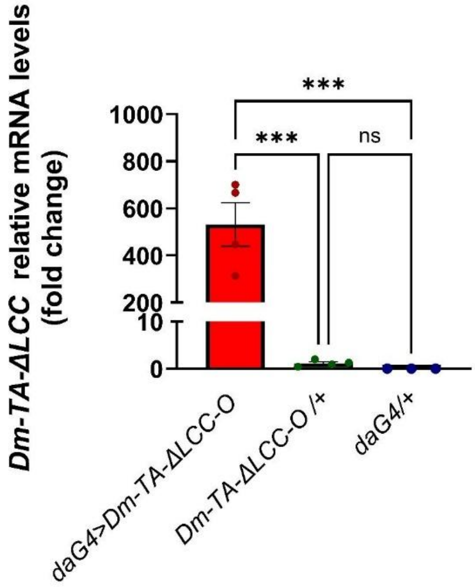


**Supplementary Figure 11.** ***Dm-TA-ΔLCC*** **mRNA expression levels (mean ± SEM) in guts of** ***daG4>Dm-TA-ΔLCC-O*** **3rd instar larvae and negative controls (*daG4/+*, and *Dm-TA-ΔLCC-O/+*).** Three-four biological replicates (dots), each with 20 guts, per genotype were analysed. *** indicates P< 0.001, significant differences in the relative expression levels determined by one-way ANOVA ($F_{2,8}$ = 28.22, P = 0.0002) and Holm-Šídák's multiple comparisons test. ns, not significant.

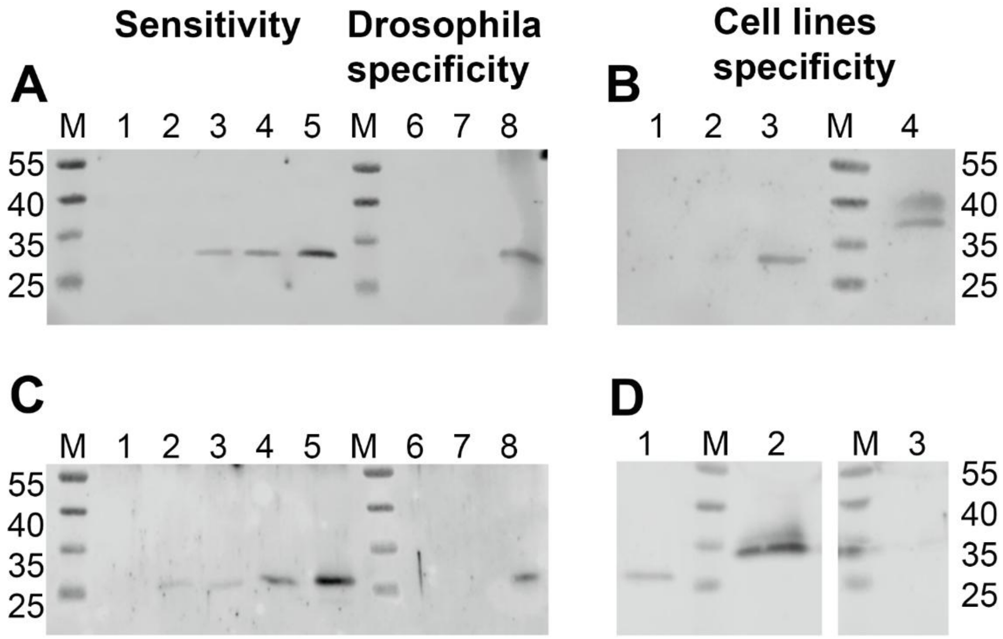


**Supplementary Figure 12. Validation of custom rabbit polyclonal antibodies anti TS-ΔIsPET and TA-ΔLCC. A** Sensitivity and specificity of the TS-ΔIsPET antibody on *D. melanogaster* extracts. Lanes 1-5: increasing amounts (10, 25, 50, 100, 200 ng) of recombinant TS-ΔIsPET protein; lanes 6 and 7: 20 and 60 µg total proteins from *D. melanogaster* negative control ($w^{1118}$) larval extracts; lane 8: 60 µg $w^{1118}$ larval extract added with 100 ng of recombinant TS-ΔIsPET protein. **B** Specificity of the TS-ΔIsPET antibody on cell culture supernatants. Lane: 1: supernatants from pAc5-STABLE-neo vector-transfected S2R+ cells (negative control); lane 2: 5 ng recombinant TS-ΔIsPET; lane 2: 10 ng recombinant TS-ΔIsPET; lane 3: supernatants of *Dm-TS-ΔIsPET*-transfected S2R+ cells at 2 days post-transfection. **C** Sensitivity and specificity of the TA-ΔLCC antibody on *D. melanogaster* extracts. Lanes 1-5: increasing amounts (10, 25, 50, 100, 200 ng) of recombinant TA-ΔLCC; lanes 6 and 7: 20 and 60 µg total proteins from *D. melanogaster* negative control ($w^{1118}$) larval extracts; lane 8: 60 µg $w^{1118}$ larval extract added with 100 ng recombinant TA-ΔLCC. **D** Specificity of the TA-ΔLCC antibody on cell culture supernatants. Lane 1: 10 ng recombinant TA-ΔLCC, lane 2: supernatants of *Dm-TA-ΔLCC*-transfected S2R+ cells at 2 days post-transfection. lane 3: supernatants from pAc5-STABLE-neo vector-transfected S2R+ cells (negative control) cell line.
Both antibodies specifically recognised their respective target proteins, with a sensitivity of 50 and 25 ng for TS-ΔIsPET and TA-ΔLCC, respectively. Non-specific signals were absent in $w^{1118}$ larval extracts or in supernatants from control cells not expressing the recombinant proteins.

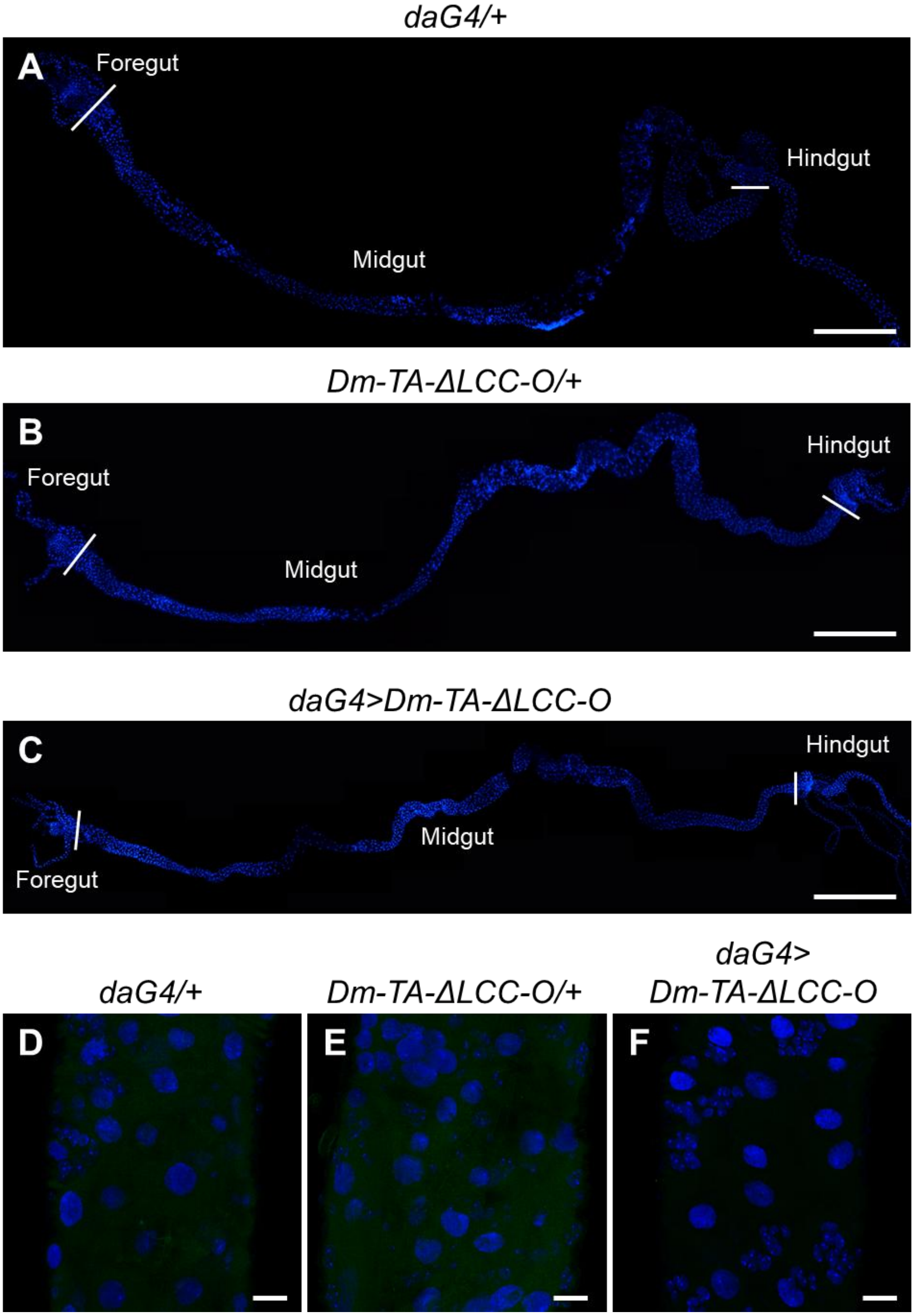


**Supplementary Figure 13. Negative controls of immunostaining experiments.** *Dm*-TA-ΔLCC whole-mount immunostaining negative controls performed on *daG4/+* (**A, D**), *Dm-TA-ΔLCC-O/+* (**B, E**), and *daG4>Dm-TA-ΔLCC-O* (**C, F**) gut samples, in which the primary antibody was omitted. Bars: 1 mm. (**A-C**), 20 µm (**D-F**).

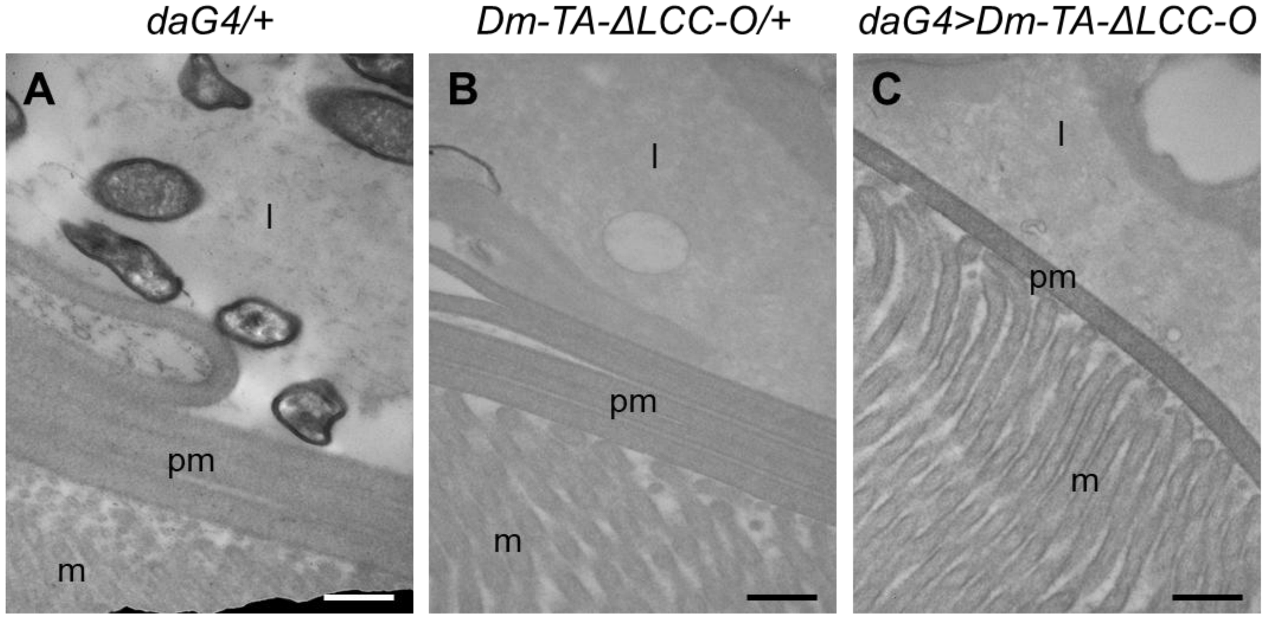


**Supplementary Figure 14. Negative controls of immunogold experiments.** *Dm*-TA-ΔLCC immunogold negative controls performed on *daG4/+* (**A**), *Dm-TA-ΔLCC-O/+* (**B**), and *daG4>Dm-TA-ΔLCC-O* (**C**) gut samples, in which the primary antibody was omitted. m: microvilli; l: lumen; pm: peritrophic matrix. Bars: 500 nm.

**Supplementary Table 1: List of primers used for qPCR analyses.**

| **Primer name** | **Sequence (5'-> 3')** |
|---|---|
| Is-PET For | TTGGAAAAAAGGGCGTGG |
| Is-PET Rev | TGGTGATGATGGTGGTGAG |
| LCC For | TAGCCGAGCAAAATCCGAG |
| LCC Rev | TGTCGTTATCCACCCAGAG |
| Rp49 For | CTAAGCTGTCGCACAAATGG |
| Rp49 Rev | TAAACGCGGTTCTGCATGAG |
